# A new genome allows the identification of genes associated with natural variation in aluminium tolerance in *Brachiaria* grasses

**DOI:** 10.1101/843870

**Authors:** Margaret Worthington, Juan Guillermo Perez, Saule Mussurova, Alexander Silva-Cordoba, Valheria Castiblanco, Charlotte Jones, Narcis Fernandez-Fuentes, Leif Skot, Sarah Dyer, Joe Tohme, Federica Di Palma, Jacobo Arango, Ian Armstead, Jose J De Vega

**Author notes:** Correspondence to; Earlham Institute, Norwich Research Park, Norwich, NR4 7UZ, UK. Department of Horticulture, University of Arkansas, 306 Plant Sciences Bldg, Fayetteville, AR, 72701, USA. NIAB, Huntingdon Road, Cambridge, CB3 0LE, UK.

## Abstract

Toxic concentrations of aluminium cations and low phosphorus availability are the main yield-limiting factors in acidic soils, which represent half of the potentially available arable land. *Brachiaria* grasses, which are commonly sown as a forage in the tropics because of their resilience and low demand for nutrients, have a greater tolerance to high concentrations of aluminium cations than most other grass crops. In this work, we explored the natural variation in tolerance to aluminium cations (Al^3+^) between high and low tolerant *Brachiaria* species and characterised their transcriptional differences during stress. We also identified three QTLs associated with root vigour during Al^3+^ stress in their hybrid progeny. By integrating these results with a new *Brachiaria* reference genome, we have identified 30 genes responsible for Al^3+^ tolerance in *Brachiaria*. We also observed differential expression during stress of genes involved in RNA translation, response signalling, cell wall composition and vesicle location genes homologous to aluminium-induced proteins involved in limiting uptake or localizing the toxin. However, there was limited regulation of malate transporters in *Brachiaria*, which are associated with external tolerance mechanisms to Al^3+^ stress in other grasses. The contrasting regulation of RNA translation and response signalling suggests response phasing is critical to Al^3+^ tolerance.

**HIGHLIGHT:** We identified QTLs, genes and molecular responses in high and low tolerant *Brachiaria* grasses associated with aspects of response to aluminium stress, such as regulation, cell-wall composition and active transport.

## INTRODUCTION

Acidic soils constitute approximately 30 % of the world’s total land area and 50 % of the potentially available arable land (Von Uexküll and Mutert, 1995). Acidic soils are particularly predominant in a northern “temperate belt” and a southern “subtropical belt”. Therefore, a broad range of vegetable, cereal and forage crops can be yield-limited in these conditions (Hede *et al.*, 2001). The adverse effects of soil acidity are mostly associated with several mineral toxicities and deficiencies, particularly increased concentrations of soluble forms of manganese, iron and aluminium, and reduced levels of available forms of phosphorus, calcium, magnesium and potassium. Among these, lower phosphorus solubility and aluminium toxicity are considered the main limiting factors on productivity (Eswaran *et al.*, 1997). As soil pH decreases below 5, aluminium becomes soluble as the aluminium trivalent cation (Al^3+^), a form highly toxic to plants. Soluble Al^3+^ effect on root apices results in diminished ion and water uptake.

Although acid soils can be conditioned for improved agricultural use through the addition of lime, in general, the most sustainable strategy is a combination of agronomic practices and growing tolerant cultivars. Natural variation in aluminium tolerance has been identified for a number of crops with rice (*Oryza sativa*) being the most aluminium-tolerant among the food staples (Rao *et al.*, 2016). All *Brachiaria* (Trin.) Griseb. (syn. *Urochloa* P. Beauv.) species show greater tolerance to Al^3+^ toxicity than most other grass crops, including maize (*Zea mays*), rice or wheat (*Triticum aestivum*) (Kochian *et al.*, 2015).

*Brachiaria* grasses are native to East Africa and are widely sown as a forage to feed ruminants across the tropics, particularly in areas with marginal soils. *Brachiaria* has a set of desirable genetic characteristics linked to drought and waterlogging tolerance, poor and acidic soils tolerance, and resistance to major diseases. However, *Brachiaria* resilience is principally a result of low demand for soil nutrients (Pizarro *et al.*, 2013). As a consequence, toxic cation levels (and not reduced levels of mineral solutes) are the limiting factors on *Brachiaria* species productivity in acidic soil conditions. Al^3+^ tolerance has been established to be a multigenic trait, though major genes can also be important (Kochian *et al.*, 2015; Ryan *et al.*, 2009). The incomplete transfer of tolerance from parents to near-isogenic lines in sorghum (*Sorghum bicolor*) (Melo *et al.*, 2013), maize (Guimaraes *et al.*, 2014), and wheat (Johnson *et al.*, 1997; Tang *et al.*, 2002) supports polygenic inheritance of Al^3+^ resistance.

Al^3+^ tolerance mechanisms are classified as external and internal tolerance mechanisms (Furlan *et al.*, 2018; Kochian *et al.*, 2015). In the first group, plants prevent Al^3+^ uptake by raising the pH in the rhizosphere or forming binding complexes by exudation of citrate, malate and oxalate. In the second group, Al^3+^ is absorbed and localized to cell organelles or the apoplast. In the tolerant rice variety Nipponbare, both exclusion mechanisms and internal detoxification are important in withstanding toxicity, while primarily internal detoxification occurs in the sensitive rice variety Modan (Roselló *et al.*, 2015).

Little is known about the genetic basis of Al^3+^ tolerance in *Brachiaria*. Nevertheless, screening experiments have evidenced different but consistent aluminium tolerance among *Brachiaria* species and cultivars, which indicates the genetic basis of the trait (Arroyave *et al.*, 2013). The three most important commercial species, *Brachiaria brizantha* (A.Rich.) Stapf., *B. decumbens* Stapf., and *B. humidicola* (Rendle) Schweick exist primarily as apomicts with varying levels of polyploidy (Valle and Savidan, 1996). The diploid sexual species *B. ruziziensis* (Germ. & C.M. Evrard) is also used in breeding as a bridge between apomictic species (Miles, 2007). *B. decumbens* is significantly more tolerant to Al^3+^ than *B. ruziziensis* (Arroyave *et al.*, 2011; Bitencourt *et al.*, 2011).

*Brachiaria* is one of the most widely used and promising forage genera in the American and African tropics. For the potential of this species to be realised, it is important that varieties are tailored to the particular demands of each environment in which it is grown (Bailey-Serres *et al.*, 2019). While genomic approaches can accelerate genetic gain from crop breeding, these approaches rely on resources that are costly and often scarce in “orphan crops”. As part of this project, we have also consequently produced some of the fundamental genomic resources previously absent for *Brachiaria*, uniquely a reference genome. We have applied these new genomic resources to develop a better understanding of the molecular basis of a key trait affecting *Brachiaria* productivity, namely Al^3+^ tolerance.

## METHODS

### Plant materials and phenotyping

The interspecific mapping population used in this work consisted of 169 genotypes of F1 progeny from a cross between the synthetic autopolyploid *B. ruziziensis* accession BRX 44-02 and the segmental allopolyploid *B. decumbens* accession CIAT 606 (cv. Basilisk). This population was generated initially to create saturate linkage maps and identify markers linked to apomixis (Worthington *et al.*, 2016). Accessions were phenotyped for aluminium tolerance at CIAT in Cali, Colombia, following Wenzl *et al*. (2006). Briefly, the experiment consisted of six cycles (replications over time) with 1-7 replicate plants per cycle. Plants were grown for 20 days in hydroponic solutions with 0 or 200 μM AlCl_3_ and phenotyped in each condition (control -C- and Al^3^+ stress -A-) for cumulative root length (RLC and RLA), root biomass (RBC and RBA), and root tip diameter (RDC and RDA) at the end of each experimental cycle. Data were transformed to meet the assumption of normality; root length and biomass were square-root transformed and root tip diameter was natural log-transformed. The PROC MIXED method (SAS v. 9.2, Cary, NC) was used to fit a totally random effect model. Genotypic best linear unbiased predictors (BLUPs) were calculated from stress (200 μM AlCl_3_) and control (0 μM AlCl_3_) conditions individually and back transformed to calculate relative root length (RRL), relative root biomass (RRB), and relative root tip diameter (RRD) ratios.

### Genome sequencing, assembly and annotation

We selected the *B. ruziziensis* genotype CIAT 26162 (2n = 2x = 18) as the source of genomic DNA. This is a semi-erect diploid accession from Burundi (−3.1167, 30.1333) that is believed to have been mutagenised with colchicine to produce the synthetic autotetraploid *B. ruziziensis* BRX 44-02, one of the progenitors of the interspecific mapping population analysed. The ploidy of CIAT 26162 has recently been verified by cytogenetics (*P. Tomaszewska, personal comm.*). Two paired-end libraries were created and sequenced in Illumina HiSeq 2500 machines in rapid run mode by the Earlham Institute (approx. 70X) and the Yale Center for Genome Analysis (approx. 30X). Additionally, a Nextera mate-pair (MP) library with insert length of 7 Kb was sequenced to improve the scaffolding. Read quality was assessed, and contaminants and adaptors removed. Illumina Nextera MP reads were required to include a fragment of the adaptor to be used in the following steps (Leggett *et al.*, 2013a). The pair-end shotgun libraries were assembled and later scaffolded using the MP library with Platanus v1.2.117, which is optimized for heterozygous genomes (Kajitani *et al.*, 2014). We did not use Platanus’ gap-closing step. Approximately 1 million Pacbio reads from this same genotype were generated in a PacBio RSII sequencer and used for gap filling using PBJelly v.15.8.24 (English *et al.*, 2012). Scaffolds shorter than 1 Kbp were filtered out. We used 31mer spectra analysis to compare the assemblies produced by different pipelines, as well as our final assembly with the assemblies from preceding steps. A K-mer spectrum is a representation of how many fixed-length words or K-mers (y-axis) appear a certain number of times or coverage (x-axis). The K-mer counting was performed with KAT (Mapleson *et al.*, 2016). The completeness of the assembly was checked with BUSCO (Simão *et al.*, 2015).

Repetitive and low complexity regions of the scaffolds were masked using RepeatMasker (Tarailo-Graovac and Chen, 2009) based on self-alignments and homology with the RepBase public database and specific databases built with RepeatModeler (Smit and Hubley, 2008). Long terminal repeat (LTR) retrotransposons were detected by LTRharvest (Ellinghaus *et al.*, 2008) and classified with RepeatClassifier. The 5′ and 3′ ends of each LTR were aligned to each other with MUSCLE (Edgar, 2004) and used to calculate the nucleotide divergence rate with the Kimura-2 parameter using MEGA6 (Tamura *et al.*, 2013). The insertion time was estimated by assuming an average substitution rate of 1.3 × 10^−8^ (Schmutz *et al.*, 2014)

Our annotation pipeline (De Vega *et al.*, 2015) uses four sources of evidence: (a) *De novo* and genome-guided *ab initio* transcripts deduced from RNA-Seq reads from the *B. ruziziensis* genotype assembled with Trinity (Grabherr *et al.*, 2011) and Tophat and Cufflinks (Trapnell *et al.*, 2012), (b) gene models predicted by Augustus (Stanke *et al.*, 2006), (c) homology-based alignments of transcripts and proteins from four close species with Exonerate and GMAP (Wu *et al.*, 2016), and (d) the repeats annotation. Finally, MIKADO (Venturini *et al.*, 2018) built the gene models to be compatible with this previous information. Proteins were compared with the NCBI non-redundant proteins and EBI’s InterPro databases and the results were imported into Blast2GO (Conesa *et al.*, 2005) to annotate the GO and GO slim terms, enzymatic protein codes and KEGG pathways. Proteins were also functionally annotated with the GO terms of any significant orthologous protein in the eggNOG database (Powell *et al.*, 2014), using the eggNOG-mapper pipeline (Huerta-Cepas *et al.*, 2017).

### Comparative genomics

Syntenic blocks between *B. ruziziensis* and foxtail millet (*Setaria italica* (L.) P. Beauv) whole genomes were identified with Minimap (Li, 2016), and plotted with D-GENIES (Cabanettes and Klopp, 2018). Previously, we filtered out any scaffold shorter than 10 Kbp that did not contain any gene. The assembly was anchored to foxtail millet by assigning each scaffold to the chromosome position where it had the longest alignment chain after combining proximal alignments. For clustering, proteins from five related species were assigned to eggNOG orthologous groups as before. A phylogenetic tree based on these data was built with MUSCLE by aligning the orthologous proteins within each eggnog cluster, filtering sets with one member per specie (*P. virgatum* was excluded from this analysis), and finally estimating the nucleotide divergence rate using MEGA v6, as before.

### Population genotyping and genetic map construction

Genotyping-by-sequencing libraries were prepared and sequenced for the 169 F1 progenies and the two parents as described in Worthington *et al.* (2016). Reads were demultiplexed according to the forward and reverse barcodes used during library preparation with FastGBS (Torkamaneh *et al.*, 2017) and adaptors and enzymatic motifs removed with Cutadapt (Martin, 2011). Reads were aligned to the genome using BWA MEM (Li, 2013). SNP calling was done for each sample with GATK’s Haplotycaller (Van der Auwera *et al.*, 2013) without the duplicated read filter (-drf DuplicateRead) and recalled for the population with GATK’s GenotypeGVCFs, in both tools with “--ploidy 4”. In agreement with filtering criteria previously tested in tetraploid *Brachiaria* samples (Worthington *et al.*, 2016), we used SNPs only, and required at least 12 reads to call a homozygous site in any sample and a minimum allele frequency of 5 %. We removed any site not called in a progenitor or in more than 20 % of the progeny. Markers that were heterozygous in only one parent and had a segregation ratio of a heterozygote to homozygote progeny of approximately 1:1 (between 0.5 and 1.75) were classified as single-dose allele (SDA) markers and used in the linkage map construction. Separate genetic linkage maps of BRX 44-02 and CIAT 606 were constructed in JoinMap v5 (Van Ooijen, 2011) using a threshold linkage logarithm of odds (LOD) score of 10 to establish linkage groups. Marker order was then determined using regression mapping with default settings. Downstream QTL analysis with MapQTL 6 (Van Ooijen and Kyazma, 2009) was conducted using only the *B. decumbens* CIAT 606 map.

### RNA-seq sequencing and analysis

Two replicate plants of *B. decumbens* CIAT 606 (cv. Basilisk) and *B. ruziziensis* BRX 44-02 were grown in high aluminium (200 μM AlCl_3_) and control (0 μM AlCl_3_) conditions in the greenhouse, as described previously. After three days of growth, roots and leaves were harvested from each plant. RNA extraction was performed with RNeasy Plant Mini kit (Qiagen, CA, USA) and sent to the sequencing service provider (Earlham Institute, Norwich, UK) where Illumina RNA-seq libraries were prepared and sequenced using the HiSeq 2500 platform. Sixteen libraries were independently generated and sequenced: two tissues (roots and leaves), from two species (*B. ruziziensis* and *B. decumbens*), in two conditions (0 and 200 μM AlCl_3_ hydroponic solutions), and two replicates. Contaminations in the raw data were discarded with Kontaminant (Leggett *et al.*, 2013b). Adaptors were removed with Cutadapt (Martin, 2011) and quality checked with FastQC. Reads were mapped to the assembled genome using STAR (Dobin *et al.*, 2013) and the gene models annotation for guidance. Counts were estimated with Stringtie (Pertea *et al.*, 2015). We used DEseq v2 (Love *et al.*, 2014) for analysing differential expression. Enriched GO terms and other categories in each group of differentially expressed genes were identified in R using TOPGO (Alexa and Rahnenfuhrer, 2010) using a Fisher’s test (FDR<0.05) and the “weight01” algorithm. The relation among GO terms was plotted in R using ggplot (Wickham and Chang, 2008). We also reanalysed using this pipeline the publicly available data (PRJNA314352) from Salgado *et al*. (2017), obtained from harvested root tips from 12 day-old *B. decumbens* cv. Basilisk seedlings screened in similar experimental conditions to those described in our study but after 8 hours of treatment (instead of 72 hours).

## RESULTS

### Root morphology in *Brachiaria* species in different aluminium cation concentrations

We demonstrated the superior aluminium tolerance of the *B. decumbens* accession CIAT 606 compared with *B. ruziziensis* accession BRX 44-02 (Figure 1). The root morphology of CIAT 606 was less affected than BRX 44-02 after growing for 20 days in high aluminium (200 μM AlCl_3_) and control (0 μM AlCl_3_) concentration hydroponic solutions. Under stress conditions, the roots of *B. decumbens* were over three times as long and had twice the biomass of *B. ruziziensis.* However, the root tip diameter increased in stress conditions in a similar ratio in both species (Supplementary table S1).

**Figure 1:**
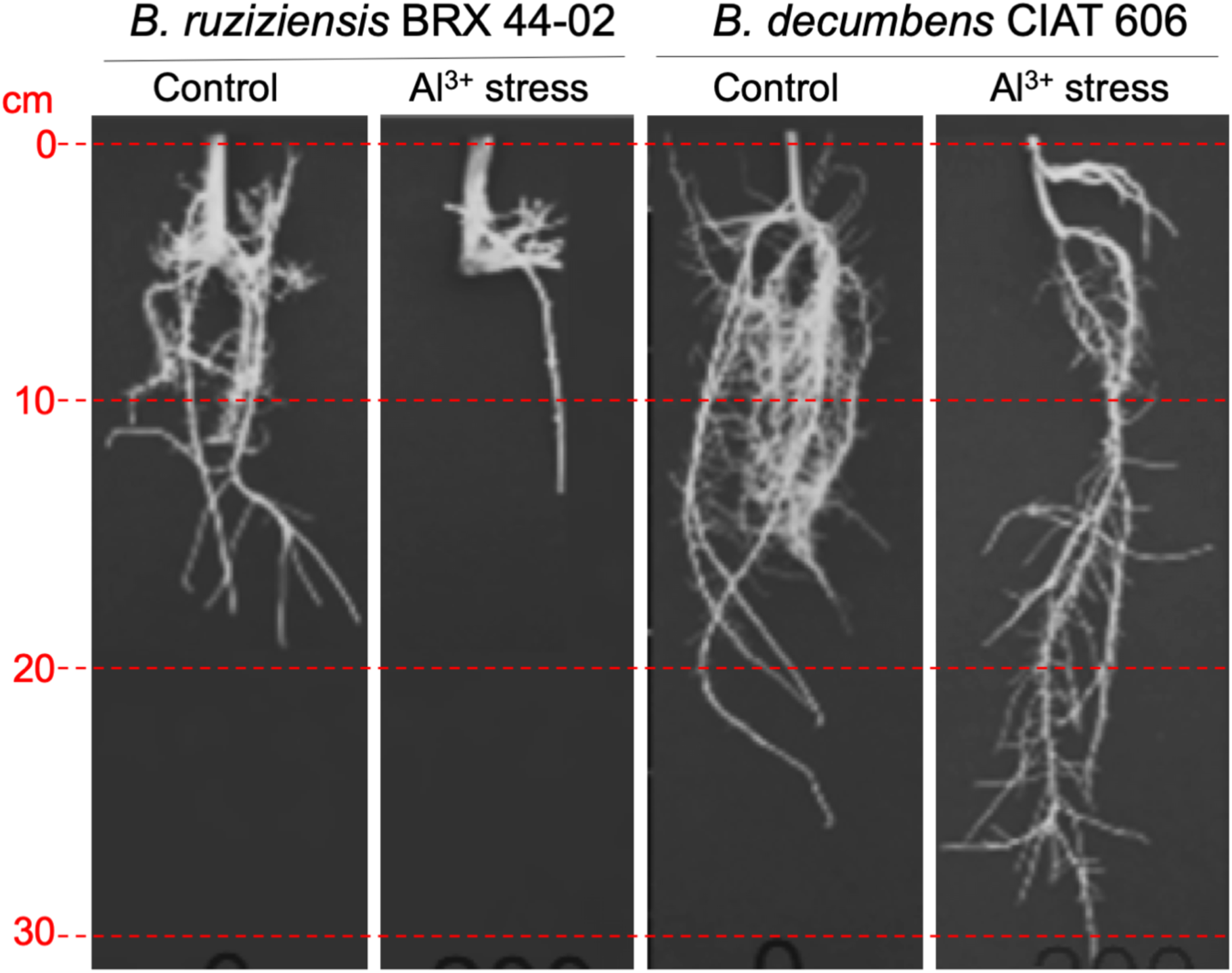
Root growth in *B. decumbens* accession CIAT 606 and *B. ruziziensis* accession BRX 44-02 (cv. Basilisk) after growing for 20 days in control and high (200 μM) Al^3+^ concentration hydroponic solutions.

The interspecific progeny obtained by crossing aluminium-tolerant *B. decumbens* accession CIAT 606 and aluminium-sensitive *B. ruziziensis* accession BRX 44-02 showed segregation in cumulative root length (RL), root biomass (RB), and root tip diameter (RD) (Figure 2; Supplementary File 1). We obtained highly significant (p < 0.001) genotypic differences in control and stress conditions for the three traits (RL, RB and RD). The root biomass (*r* = 0.73), length (*r* = 0.76) and diameter (*r* = 0.69) of the progeny were significantly correlated when grown in stress (200 μM AlCl_3_) and non-stress (0 μM AlCl_3_) conditions.

**Figure 2:**
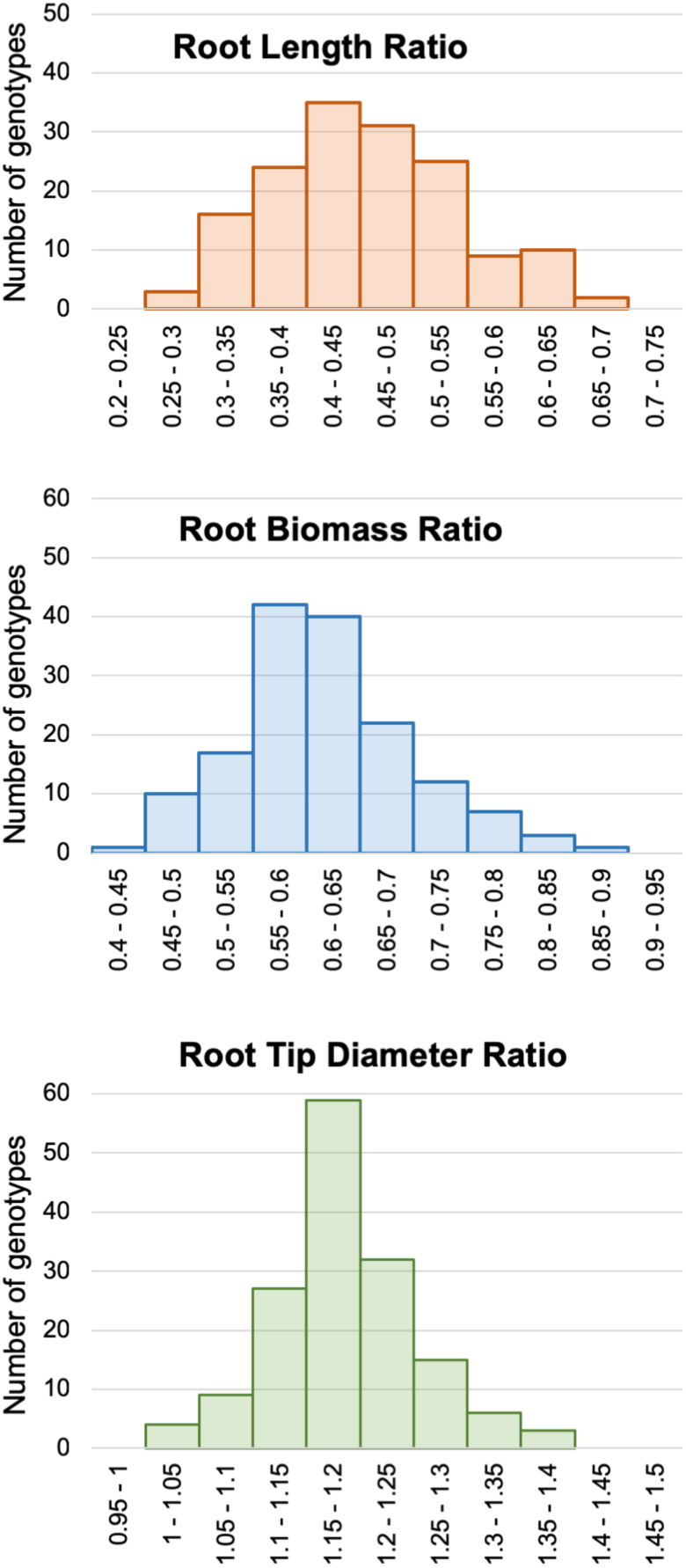
Root length (RL), root biomass (RB), and root tip diameter (RD) ratio values between Al^3+^ stress and control measured values (stress/control) in the interspecific progeny (n=169) between *B. ruziziensis* BRX 44-02 and *B. decumbens* CIAT 606 (cv. Basilisk).

### Assembly and annotation of a *Brachiaria* reference genome

A whole-genome assembly (WGA) of CIAT 26162 was produced using Platanus v.1.2.4 (Kajitani *et al.*, 2014), from Illumina paired-end and Nextera mate-pair reads with a coverage of approximately 100X and 7X, respectively (Supplementary table S2). Platanus outperformed the contiguity results obtained with other pipelines. Although the combination of ABySS for the contigs assembly and SOAP2 for the scaffolding resulted in a larger assembly, a kmer frequency analysis (Mapleson *et al.*, 2016) evidenced that the additional content was repeated under-collapsed heterozygosity that Platanus had purged instead (Supplementary figure S1). A gap-filling step using approximately 1 million Pacbio reads with an average length of 4.8 Kbp resulted in a reduced percentage of ambiguous nucleotides (Ns) from 17.45 % to 11.39 %. We finally discarded all the sequences under 1 Kbp to produce the reference genome we used for the downstream analysis (Table 1). To assess the completeness of the assembly, we verified that 1,345 (93.4 %) of the 1,440 BUSCO orthologous genes (Simão *et al.*, 2015) were present in the assembly; 1,216 of which were completed and in a single copy, 32 were duplicated, and 97 were fragmented. This WGA is deposited at NCBI with the accession number GCA_003016355. The raw reads are deposited in the Bioproject PRJNA437375.

**Table 1:**
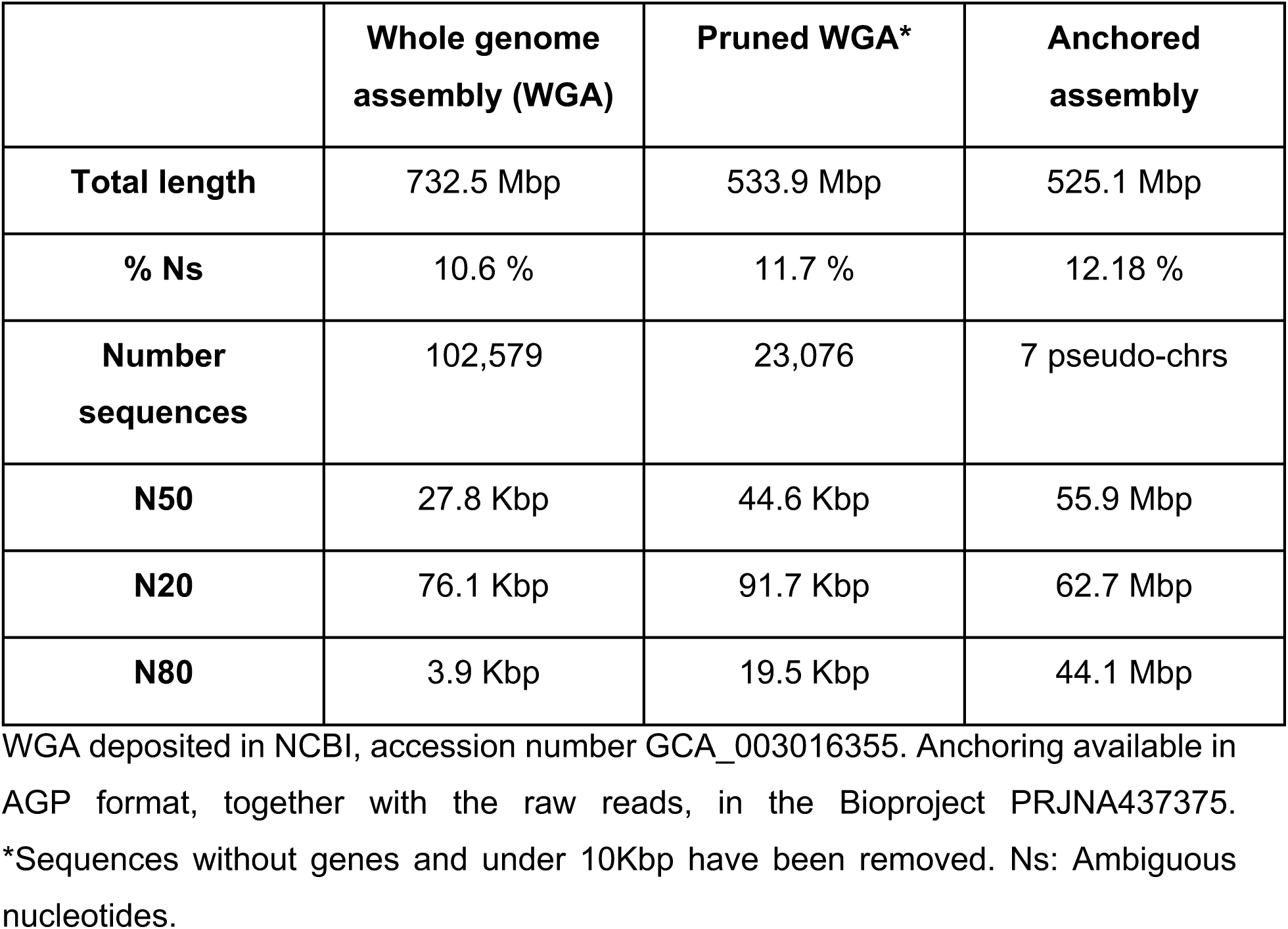
Statistics associated with the assembly of the *Brachiaria ruziziensis* reference genome and anchored in pseudo-molecules.

|  | **Whole genome**<br>**assembly (WGA)** | **Pruned WGA\*** | **Anchored**<br>**assembly** |
| --- | --- | --- | --- |
| **Total length** | 732.5 Mbp | 533.9 Mbp | 525.1 Mbp |
| **% Ns** | 10.6 % | 11.7 % | 12.18 % |
| **Number**<br>**sequences** | 102,579 | 23,076 | 7 pseudo-chrs |
| **N50** | 27.8 Kbp | 44.6 Kbp | 55.9 Mbp |
| **N20** | 76.1 Kbp | 91.7 Kbp | 62.7 Mbp |
| **N80** | 3.9 Kbp | 19.5 Kbp | 44.1 Mbp |

WGA deposited in NCBI, accession number GCA\_003016355. Anchoring available in AGP format, together with the raw reads, in the Bioproject PRJNA437375. \*Sequences without genes and under 10Kbp have been removed. Ns: Ambiguous nucleotides.

**Table 2:** *Brachiaria* genes differentially expressed during Al^3+^ in *Brachiaria* highlighted in our analysis.

| GeneID | Bruz | Bdec | Gene name | *P. halli* | *S. italica* | *S. viridis* | *A. thaliana* | Uniprot | Function and induction |
| --- | --- | --- | --- | --- | --- | --- | --- | --- | --- |
| 29G2 | 0.71 | 0.17 | STOP1 | 3G044400 | 7G287100 | 7G299300 | AT1G34370 | STOP1_ORYSJ | Zinc-finger Transcription factor involved in aluminium tolerance |
| 61G2 | 0.88 | 0.23 | Citrate transporter detoxification 42 | 9G552400 | 9G489200 | 9G493200 | AT1G51340 | DTX42_ARATH | Citrate transporter responsible for citrate exudation into the rhizosphere to protect roots from aluminium toxicity. Up-regulated by aluminium. Expressed in roots, but not in shoots. |
| 94G8 | 7.68 | 2.36 | P transmembrane transporter | 7G133200 | 7G082600 | 7G090100 | AT4G13420 | HAK1_ORYSJ | **In the QTL.** Potassium transmembrane transported induced by P starvation exclusively in roots. |
| 126G26 | 1.83 | 0.95 | STOP1 | 6G306100 | 6G252000 | 6G256600 | AT5G22890 | STOP1_ORYSJ | Transcription factor involved in aluminium tolerance |
| 259G14 | 2.36 | 1.43 | Citrate transporter protein detox. 42 | 9G201000 | 9G204700 | 9G203100 | AT1G51340 | DTX42_ARATH | Citrate transporter responsible for citrate exudation into the rhizosphere to protect roots from aluminium toxicity. Up-regulated by aluminium. Expressed in roots, but not in shoots. |
| 462G24 | -2.35 | 0.38 | AMLT10 | 4G147500 | 4G162600 | 4G130000 | - | K3XVN8_SETIT | Aluminium-activated malate transporter 10-like |
| 598G8 | -0.97 | -2.58 | Cytochrome P450 | 7G003400 | 1G019400 | 1G019300 | AT5G07990 | C93G1_ORYS | **In the QTL.** Metal-binding transmembrane P450 cytochromes |
| 675G12 | 4.27 | 3.89 | Citrate transporter protein detox. 42 | 5G042400 | 9G489200 | 9G493200 | AT1G51340 | DTX42_ARATH | Citrate transporter responsible for citrate exudation into the rhizosphere to protect roots from aluminium toxicity. Up-regulated by aluminium. Expressed in roots, but not in shoots. |
| 869G16 | 1.36 | 1.83 | Cysteine XCP1 protease | 2G435100 | 8G204200 | 8G213900 | AT4G35350 | XCP1_ARATH | **In the QTL.** A proteinase that may have a developmental senescence specific cell death function during apoptosis and metal detoxification |
| 1615G2 | 2.69 | 1.29 | STOP1 | 6G306100 | 6G252000 | 6G256600 | AT5G22890 | STOP1_ORYSJ | Zinc-finger Transcription factor involved in aluminium tolerance |
| 1984G2 | 1.09 | 0.82 | SLK2 | 4G345700 | 4G015400 | 4G015300 | AT5G62090 | SLK2_ARATH | DNA-binding adapter subunit of the SEU-SLK2 transcriptional corepressor of abiotic stress (e.g. salt and osmotic stress) response genes by facilitating auxin response and sustaining meristematic potential. |
| 2499G8 | 1.18 | -0.15 | OsTITANIA/AtOB E3 | 9G552000 | 9G488800 | 9G492800 | AT1G14740 | TTA1_ORYSJ | Widely expressed transcription factor that functions as regulator of metal transporter genes responsible for essential metals delivery to shoots and normal plant growth. Required for the maintenance of metal transporter gene expression, such as IRT1, IRT2, ZIP1, ZIP9, NRAMP1 and NRAMP5. |
| 2708G4 | 1.64 | 1.94 | Auxin transport protein BIG | 2G178200 | 2G154800 | 2G160200 | AT3G02260 | BIG_ORYSJ | Auxin-mediated developmental responses (e.g. cell elongation, apical dominance, lateral root production, inflorescence architecture, general growth and development). |
| 3096G6 | 0.51 | 0.27 | ALS1 | 9G083700 | 9G088000 | 9G086400 | AT5G39040 | AB25B_ORYSJ | Metal transporter involved in the sequestration of aluminium into vacuoles, which is required for cellular detoxification of aluminium. Induced by aluminium in roots. Expressed in primary roots and lateral roots. |
| 3672G6 | 2.54 | 2.26 | NRAT1 | 1G025100 | 1G098100 | 1G097200 | AT1G80830 | NRAT1_ORYSJ | Metal transporter Al<sup>3+</sup>, but not divalent cations Fe<sup>2+</sup>, Mg<sup>2+</sup>, Cd<sup>2+</sup>. Involved in Al tolerance by taking up Al in root cells, where it is detoxified by chelation with organic acid anions and sequestration into the vacuoles. Induced by aluminium in roots. Positively regulated by ART1. |
| 4534G2 | 4.74 | 0.43 | SAD2 importin beta-like transporter | 7G101600 | 7G051600 | 7G057300 | AT2G31660 | SAD2_ARATH | Involved in the regulation of the abscisic acid (ABA)-mediated pathway in response to cold or salt stress, UV-B responses. Involved in trichome initiation. Negative regulator miRNA activity. Expressed in roots, epidermal and guard cells of leaves, stems and siliques |
| 5233G2 | 1.07 | 1.18 | XTH23 | 4G029300 | 4G246200 | 4G258800 | AT4G25810 | XTH23_ARATH | Xyloglucan endotransglucosylase hydrolase protein 23 cleaves and religates xyloglucan polymers, an essential constituent of the primary cell wall, and thereby participates in cell wall construction of growing tissues. Up-regulated by abscisic acid (ABA). |
| 5400G2 | -1.38 | -0.05 | XTH | 8G197800 | 8G142000 | 8G151900 | AT5G13870 | XTH5_ARATH | Cleaves and religates xyloglucan polymers, an essential constituent of the primary cell wall, and thereby participates in cell wall construction of growing tissues. Strongly down regulated by abscisic acid. Root specific. |
| 6544G2 | -0.07 | 0.95 | XTH32 | 2G372300 | 2G313800 | 2G324800 | AT2G36870 | XTH8_ORYSJ | Xyloglucan endotransglucosylase hydrolase protein 32 cleaves and religates xyloglucan polymers, an essential constituent of the primary cell wall, and thereby participates in cell wall construction of growing tissues. May be involved in cell elongation processes. Induced by gibberellic acid (GA3). |
| 6944G2 | 1.92 | 2.01 | Phospholipid transporting ATPase | 4G247400 | 4G147300 | 4G165700 | AT1G26130 | ALA12_ARATH | Involved in transport of phospholipids. |
| 7361G2 | 2.26 | 1.39 | STAR2/ALS3 | 3G099200 | 3G043600 | 3G044400 | AT2G37330 | STAR2_ORYSJ | Associates with STAR2 to form a functional transmembrane ABC transporter required for detoxification of aluminium (Al) in roots. Can specifically transport UDP-glucose. Induced by Al in roots. Positively regulated by ART1. Root specific. |
| 8068G4 | 2.77 | 2.12 | Mg transporter MRS2-D | 7G150800 | 7G099600 | 7G107500 | AT3G19640 | MRS2D_ORYSJ | Putative magnesium transporter. |
| 8234G4 | -1.40 | -1.30 | Uncharacterised | - | - | - | - | A0A3L6T2E7_PANMI | In the QTL. Uncharacterised protein. |
| 8440G2 | 0.88 | -1.68 | Cytochrome P450 | 6G227500 | 2G094000 | 2G096800 | AT4G36380 | C90C1_ARATH | In the QTL. Metal-binding transmembrane P450 cytochromes |
| 8448G4 | 0.59 | 1.26 | STAR1 | 4G029500 | 4G246800 | 4G259400 | AT1G67940 | STAR1_ORYSJ | Associates with STAR2 to form a functional transmembrane ABC transporter required for detoxification of aluminium (Al) in roots. Can specifically transport UDP-glucose. Induced by Al in roots. Positively regulated by ART1. Root specific. |
| 11634G4 | 0.04 | 4.75 | AMLT1 | 7G143300 | 7G007400 | 7G102000 | AT3G11680 | ALMT8_ARATH | Malate transporter. |
| 11683G4 | 1.00 | 0.66 | XTH27 | - | - | - | - | A0A1D6K9K3_ZM | Xyloglucan endotransglucosylase hydrolase protein 27 cleaves and religates xyloglucan polymers, an essential constituent of the primary cell wall, and thereby participates in cell wall construction of growing tissues. |
| 12087G2 | 1.31 | 3.21 | Phosphate transporter pho1-2 | 1G440700 | 1G360400 | 1G367100 | AT3G23430 | PHO12_ORYSJ | Involved in the transfer of inorganic phosphate (Pi) from roots to shoots. Not induced by Pi deficiency in roots. Specifically expressed in roots. |
| 26018G2 | 1.08 | 0.98 | transporter NRT1 nitrate transporter | 9G390400 | 9G327900 | 9G333900 | AT2G26690 | PTR27_ARATH | Low-affinity proton-dependent nitrate transporter. Up-regulated in the shoots by nitrate, but no changes in the roots. |
| 30107G2 | 3.24 | 3.02 | NRAT1 | 2G074500 | 1G098100 | 1G097200 | AT1G80830 | NRAT1_ORYSJ | Metal transporter Al<sup>3+</sup>, but not divalent cations Fe<sup>2+</sup>, Mg<sup>2+</sup>, Cd<sup>2+</sup>. Involved in Al tolerance by taking up Al in root cells, where it is detoxified by chelation with organic acid anions and sequestration into the vacuoles. Induced by aluminium in roots. Positively regulated by ART1. |

GenelD: Gene number in our reference genome. Bruz/Bdec: Differential expression fold-change values in *B. ruziziensis* and *B.* *decumbens*, significant p-values under 0.05 are coloured. *P. halli*/*S. italica*/*S. viridis*/*A. thaliana*: Top Blastp result in each of these species. Uniprot and "Function and induction": Top Blastp result and associated annotation in the Uniprot database.

The repeat content (Supplementary table S3) was 51 % of the total genome (656,355,643 bp, after excluding ambiguous bases, Ns), which is close to the 46 % repeat content in foxtail millet (*Setaria italica* (L.) P. Beauv) (Zhang *et al.*, 2012). We found a large number of *Gypsy* and *Copia* LTRs, which represent 24 % and 9.5 % of the total genome excluding Ns. These transposons and proportions are also very similar to those observed in foxtail millet. We compared the divergence between the flanking tails in the LTRs (Supplementary figure S2) and identified a single very recent burst of LTR *Gypsy* activity around 0.6 MYA (Kimura = 0.042 ± 0.026) and of LTR *Copia* also around 0.6 MYA (Kimura = 0.041 ± 0.027). Other repeat elements, including LINEs (Long Interspersed Nuclear Elements), simple repeat patterns of the sequence, satellites, and transposons were much less common, except for En/Spm DNA transposons observed in 4.2 % of the genome.

We annotated 42,232 coding genes, which included 42,359 predictive open reading frames (ORFs), as well as 875 non-coding genes without a predicted ORF (Supplementary file 2). Together these transcripts and non-coding genes define 43,234 targets for the expression analysis. 35,982 of the coding transcripts had a homologous protein in the NCBI non-redundant (*nr*) database. In 58% of the cases, the top hit was a *S. italica* sequence (Supplementary figure S3). Among those 35,982, 33,963 were functionally annotated with at least one GO term, and 39,488 had an InterPro annotation. We also identified the best reciprocal hits with *Arabidopsis thaliana*, rice, *Panicum halli* Vasey, foxtail millet, and *Setaria viridis* (L.) Beauv; and the top homologous protein in Uniprot (Supplementary file 3). We aligned the transcripts and proteins from five sequenced species in the subfamily Panicoideae, foxtail millet, *S. viridis*, maize, *P. halli* and switchgrass (*P. virgatum* L.) on the WGA, and found that on average 78 % and 72 % of the transcripts and proteins aligned with an identity over 0.7, respectively (Supplementary table S4).

### Comparative genomics with related grasses

We could assign 35,831 *Brachiaria* proteins to an eggNOG orthologous group (Powell *et al.*, 2014), and 13,570 proteins could be further annotated with GO terms from the eggNOG database. We also assigned the proteins from other species in the Poaceae family to these eggNOG orthologous groups for Poaceae (poaVIR) in the eggNOG database in order to identify shared clusters of proteins among these species (Supplementary file 4; Supplementary figure S4). More than 70 % of the clusters of proteins had double or triple the number of proteins in switchgrass than the other species because of relatively recent whole-genome duplication events (Supplementary table S5). Approximately 20 % of the clusters in maize and *B. ruziziensis* had two proteins. From this analysis, we also estimated that there are approximately two thousand proteins in other close species that were missed in our *Brachiaria* assembly.

We estimated the divergence between these species based on the Kimura divergence values between pairs of orthologous proteins in 6,450 clusters of proteins that included only one member from each species (*P. virgatum* was excluded from this analysis; Supplementary figure S5). By assuming an average substitution rate of two-times (diploid) 1.3 × 10^−8^ (Schmutz *et al.*, 2014), we estimated that *Brachiaria* diverged from the other Paniceae genera, *Setaria* and *Panicum*, around 13.4-15.5 million years ago (MYA), while the split of the Paniceae and Andropogonodae tribes of subfamily Panicoideae took place around 23.8-26.3 MYA (Supplementary figure S6). The 23,076 scaffolds in the WGS longer than 10 Kbp or with at least one annotated gene (533.9 Mbp) were aligned with the nine chromosomes of foxtail millet, the closest relative with a high quality sequenced genome (Zhang *et al.*, 2012). Up to 21,145 of the 23,076 scaffolds (91.6 %), which comprise 525.1 Mbp, could be aligned (Figure 3; Supplementary file S5). Furthermore, we assigned chromosomal positions to 41,847 coding genes (41,974 transcripts) contained in these anchored sequences (Supplementary file S6). We identified 59 synteny blocks, 36 of which were longer than 1 Mbp in both species (Figure 3). There were three large translocations when comparing the foxtail millet and *B. ruziziensis* genomes between chromosomes 1 and 7, 2 and 6, as well as 3 and 5. Four inversions (smaller than the translocations) were identified between tails in chromosomes 1 and 4 (both ends), 2 and 9 (proximal end), and 2 and 3 (proximal end).

**Figure 3:**
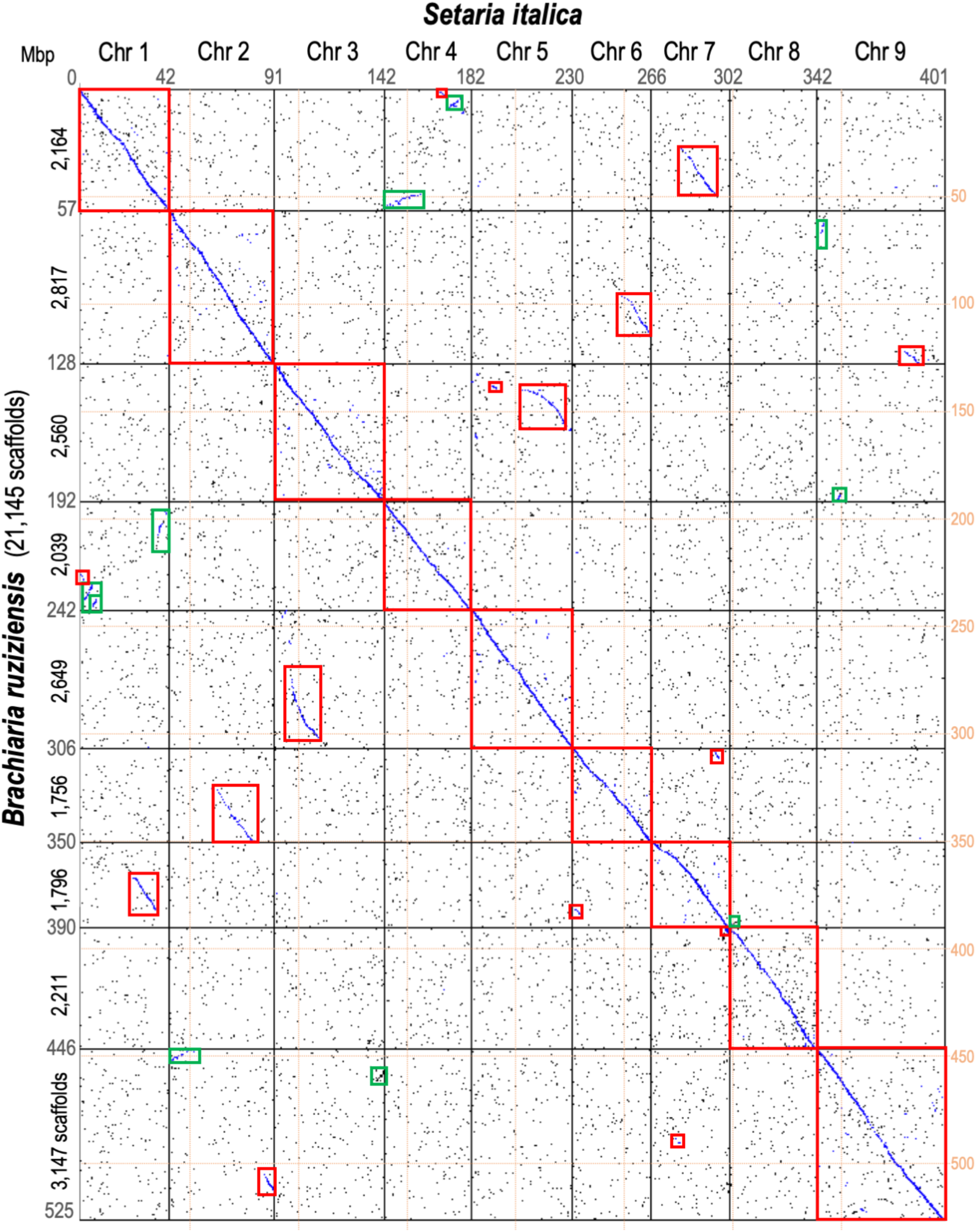
Synteny between the *Brachiaria ruziziensis* and foxtail millet (*Setaria italica*) genomes. The 36 synteny blocks longer than 1 Mbp and translocations are highlighted in red or green boxes according to their direction.

### QTL mapping in the interspecific *B. ruziziensis* X *B. decumbens* population

Between 78.8 and 91.5 % of the Genotyping-by-sequencing (GBS) reads from each 169 interspecific progeny and the parents could be aligned in the assembly. After filtering, there were an average of 81,831 SNPs and 15,595 indel sites per sample. In total, 799,155 polymorphic sites were called in the population, 85.7 % of these sites were SNPs. After filtering, 15,074 SNP sites were homozygous in the female parent (*B. ruziziensis* BRX 44-02) and heterozygous in the male parent (*B. decumbens* CIAT 606) (annotated as nnxnp markers in Joinmap), 4,891 sites were heterozygous in only the female parent (lmxll), and 1,652 were heterozygous in both parents (hkxhk). We classified 4,817 nnxnp and 1,252 lmxll sites as single dose alleles (SDAs, or “simplex” markers) based on their 1:1 heterozygous/homozygous segregation ratio in the progeny and used them in the genetic map construction. We used in our analysis the genetic map of the male parent (*B. decumbens* CIAT 606), which included 4,427 markers placed in 18 linkage groups (Supplementary figure S7, Supplementary file 7). This corresponds with the number of base chromosomes expected in an allotetraploid (2n = 4x = 36) *Brachiaria* population. Linkage groups had an average length of 74.7 ± 22.7 cM. Using the same raw data, a join genetic map of both parents and composed of 36 linkage groups was generated in Worthington *et al*. (2016). Two linkage groups were assigned to each the nine base chromosomes of *B. ruziziensis* based on alignments of each SNP to the WGA and named following the numbering system used for foxtail millet. We also aligned the WGA scaffolds with the genetic map to compare the co-linearity between the position of each marker in the genetic map and genome assembly (Supplementary file S7).

QTL mapping was performed to identify genetic regions associated with root length (RL), biomass (RB) and root tip diameter (RD) under control and Al^3+^ stress conditions in the interspecific mapping population. Three hundred and seventy-one markers had an LOD over 3 associated with one or more of the traits. Two hundred and twelve WGS scaffolds contained at least one marker with LOD over 3 (Supplementary file 7).

Three significant QTL were identified (Figure 4). The first QTL, which peaked at 5.22, 12.65, 25.797 and 26.027 cM and extended from 5.22 to 32.481 cM on LG 1 (Chr 8), was associated with root length and biomass under control and Al^3+^ stress conditions but not with relative root length or relative root biomass. The second QTL, which peaked at 79.738 and 96.802 cM and extended from 79.738 to 98.851 cM on LG 3 (Chr 7) was associated with relative root length, root tip diameter under Al^3+^ stress conditions, and relative root tip diameter. The last QTL extended from 49.12 to 62.127 cM (peak at 50.423 cM) on LG 4 (Chr 3) and was only associated with relative root tip diameter. These three QTLs each explained from 12.8 % to 16.1% of the phenotypic variance for the associated traits (Supplementary table S6).

**Figure 4:**
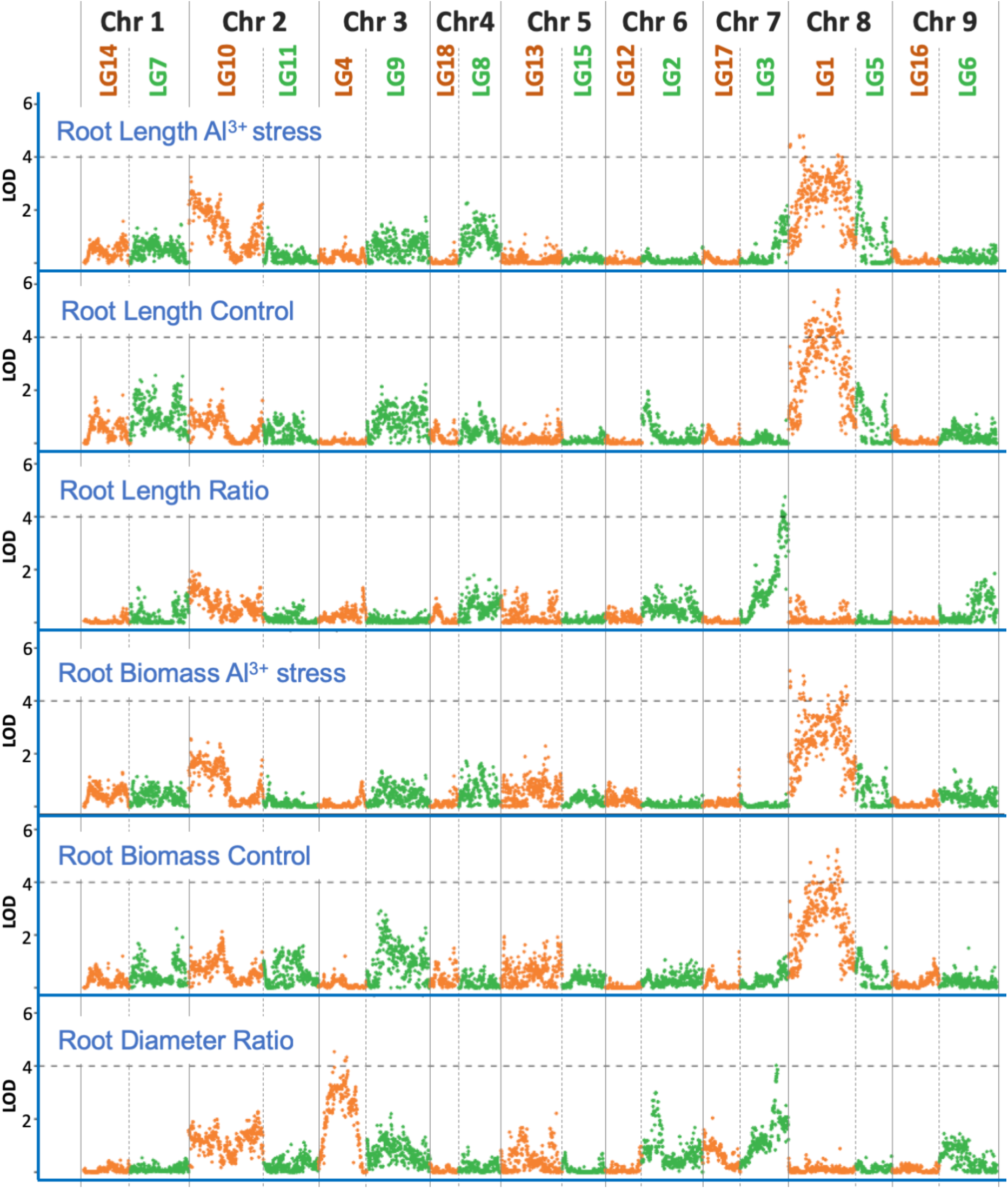
4,427 genetic markers placed in 18 linkage groups. We defined three QTLs each in LG 1 (Chr 8) for root length and biomass under control and Al^3+^ stress conditions, LG 3 (Chr 7) for root length ratio and root tip diameter ratio (stress/control), and LG 4 (Chr 3) for root tip diameter ratio.

### Transcriptional differences during stress between *Brachiaria* species

We performed RNA-seq from stem and root tissue samples extracted from *B. decumbens* CIAT 606 and *B. ruziziensis* BRX 44-02 after growing for three days in control (0 μM AlCl_3_) or high (200 μM AlCl^3^) aluminium cation concentration hydroponic solutions. When the normalised counts for all the genes were used to cluster the samples, these clusters firstly grouped by tissue, secondly by genotype, and thirdly by treatment (Supplementary figure S8). There were 4,481 differentially expressed (DE) genes in total, with 3,996 of these differentially regulated in a single genotype and tissue (Figure 5). Among these 4,481 DE genes, 4,162 genes were DE in roots only, 265 in stems only, and 54 in both tissues.

**Figure 5:**
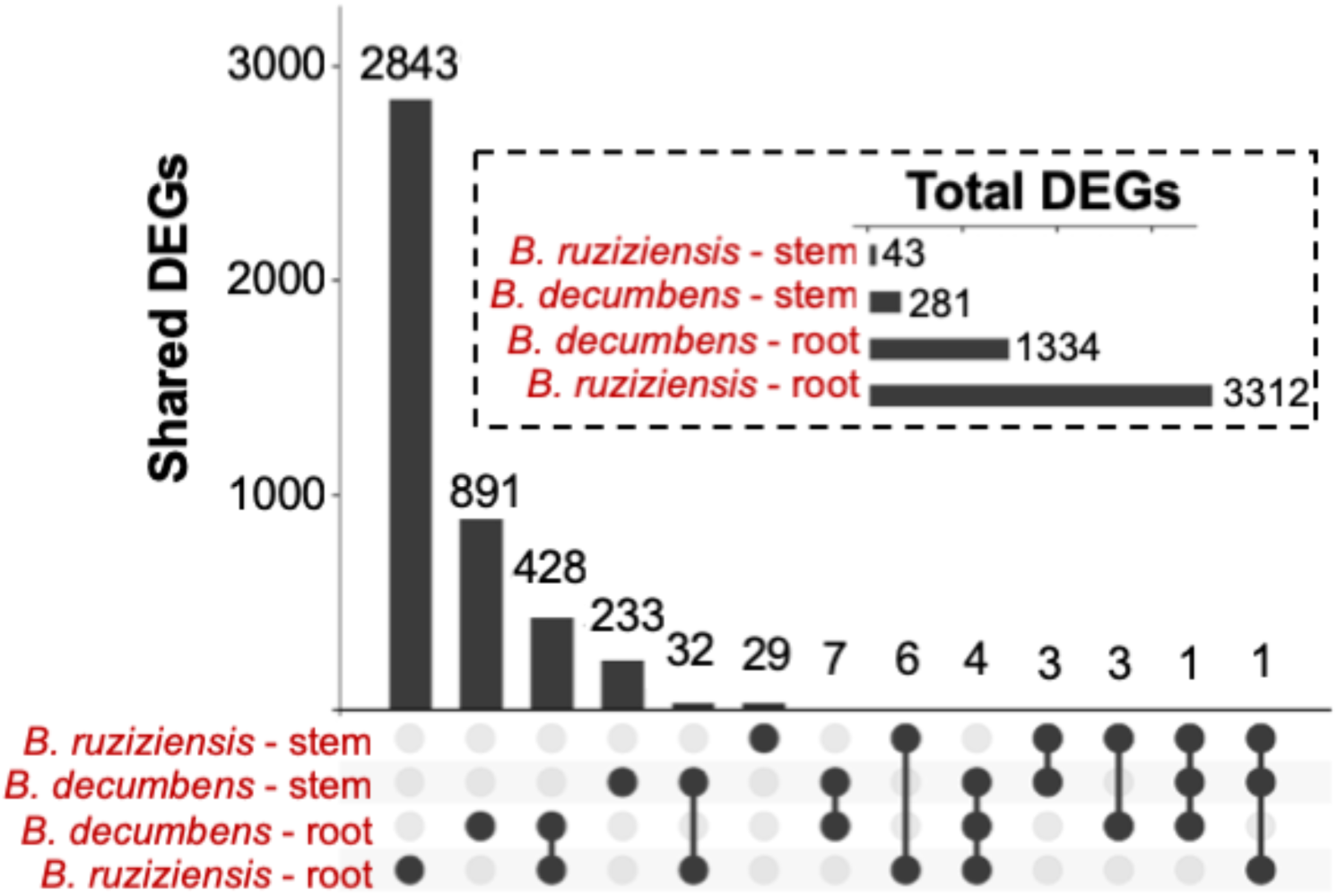
Differentially expressed (DE) genes in roots and stems from the susceptible genotype *B. ruziziensis* BRX 44-02 and the tolerant genotype *B. decumbens* CIAT 606 (cv. Basilisk) between control and high (200 μM AlCl_3_) aluminium cation concentrations in hydroponic experiments at CIAT.

The three QTL regions contained 918 genes; 581 in LG1, 153 in LG3, and 184 in LG4. Of these 918, 84 were DE genes (Supplementary file 8). Twenty-seven DE genes were annotated as membrane components, 21 were annotated as involved in response to hormones or biotic/abiotic stress, and 23 were annotated as binding to different molecular compounds, including ATP/ADP/GTP, metal ions and DNA. 34 of the 87 genes were not annotated with one of these previous GO terms, but several were annotated with several GO terms.

An enrichment analysis of GO terms over-represented among DE genes in each species allowed us to identify the biological processes (BP) and molecular functions (MF) that are similarly or differently regulated among them. After annotating the genes with the full set of GO terms (Supplementary figure S9 and S10), we also simplified the results to “GO slim” terms, contain the subset of higher-level terms from the GO resource (Figure 6; Supplementary Table S7). For the enrichment analysis we considered the either up-regulated or down-regulated genes in each sample. There was limited overlap among GO terms based on the DE genes included in each term (Supplementary figure S11).

**Figure 6:**
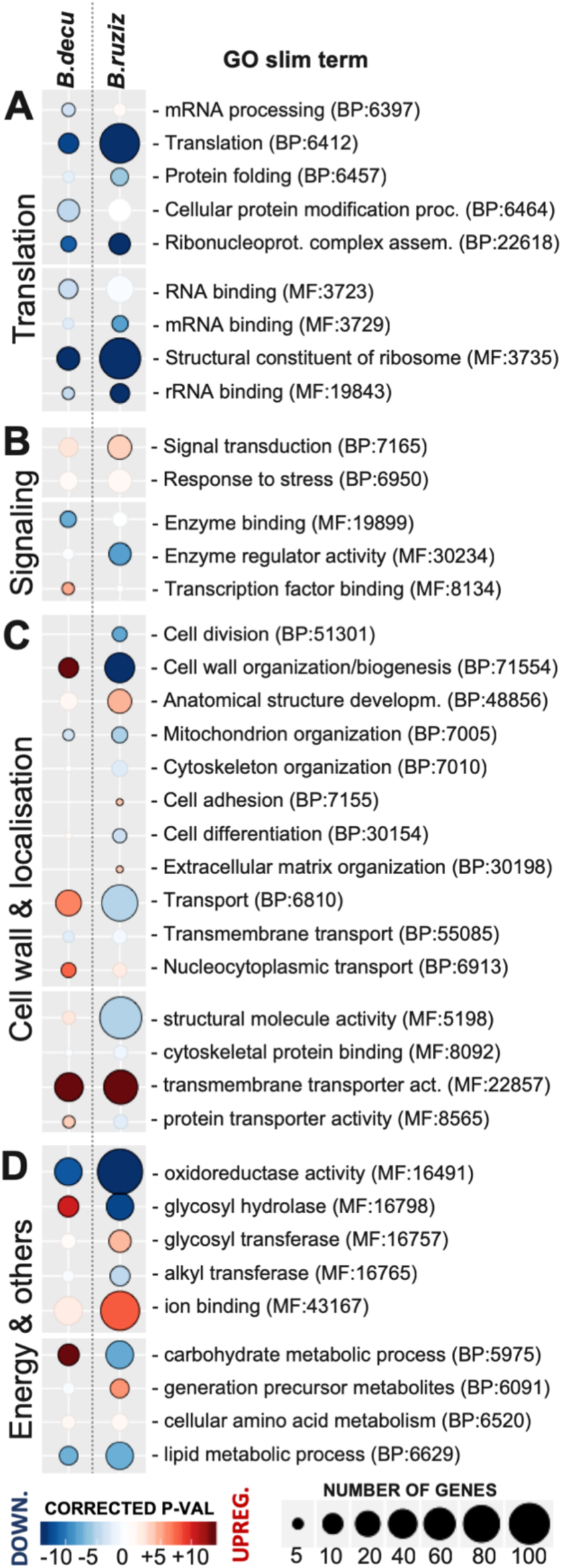
Enrichment analysis of the GO slim terms overrepresented among differentially expressed genes that are either up-regulated (red) or down-regulated (blue) during stress in roots from the susceptible *B. ruziziensis* BRX 44-02 (*B.ruziz*) or the tolerant genotype *B. decumbens* CIAT 606 cv. Basilisk (*B.decu*) genotypes.

Five MF terms related to RNA/mRNA/rRNA binding with the ribosome (MF:3723, 3729, 19843; and 3735 and 5198) and two BP terms, “translation” and “ribonucleoprotein complex assembly” (BP: 6412, 22618), were significantly enriched among down-regulated during stress genes in *B. ruziziensis* (146 genes) and *B. decumbens* (39 genes).

Genes annotated as “transmembrane transporters” (MF:22857) were highly over-represented among up-regulated during stress DE genes (44 in *B. decumbens* and 70 in *B. ruziziensis*). Fifteen DE transmembrane transporters were common to both species, and five were annotated as induced by aluminium and are later discussed.

Several GO terms showed contrasting regulation between both species. E.g. “Cell wall organization and biogenesis” was enriched in both species and included 51 down-regulated DE genes in *B. ruziziensis*, but 17 up-regulated DE genes in *B. decumbens*. Most of these genes were peroxidases and expansin proteins induced by various plant hormones (e.g. ethylene, GA, auxin) and involved in toxin removal during oxidative stress (Kochian *et al.*, 2015). The GO term “glycosyl hydrolase” was also enriched among down-regulated genes in *B. ruziziensis* (39 genes), but among up-regulated genes in *B. decumbens* (18 genes). Most of the genes annotated as “Glycosyl hydrolases” were also annotated as involved in the “carbohydrate metabolism” (BP:5975).

Several of the DE genes that exhibited high fold-changes in both species were involved in phospholipid, phosphate, magnesium, auxin, nitrate and miRNA transport in the cell (Genes 6944G2, 12087G2, 8068G4, 26018G2, 4534G2, 2708G4).

By reanalysing the data from Salgado *et al*. 2017, we could also compare the enriched GO terms between *B. decumbens* cv. Basilisk exposed to 200 μM AlCl_3_ for 8 hours and 72 hours (Supplementary Figure S12 and Supplementary Table S7). This comparison evidenced differences between 8 hours and 72 hours in the same genotype in GO terms related to RNA translation and response to stress (BP:6950), among others.

## DISCUSSION

### Forward genetics and newly produced genomic resources to identify candidate loci for abiotic stress tolerance

The negative effects of soil acidity on crop production are associated with different mineral toxicities and deficiencies. Aluminium cation toxicity has long been established to be the single most important limiting factor of acidic soil productivity (Eswaran *et al.*, 1997). All *Brachiaria* species show some tolerance to Al^3+^ toxicity, particularly compared with other grasses such as wheat, rice and maize (Arroyave *et al.*, 2013; Kochian *et al.*, 2015). While most crops reduce root growth to 50 % when exposed to 5 μM Al^3+^, *Brachiaria* species need to be exposed to up to 35 μM to exhibit a reduced root growth of 25 % (Poschenrieder *et al.*, 2008).

Natural adaptation to acid soils can be measured by estimating root vigour in low pH conditions and tolerance to high concentrations of Al^3+^ (Wenzl *et al.*, 2006). In our experiment, *B. decumbens* CIAT 606 (cv. Basilisk) root morphology was less affected than *B. ruziziensis* BRX 44-02 after growing for 20 days in control and high 200 μM Al^3+^ concentration hydroponic solutions. As a comparison, in a 21-day long term study using a similar experimental protocol, four *B. decumbens* and four *B. ruziziensis* genotypes showed intermediate tolerance of Al^3+^, one *B. ruzizensis* genotype was sensitive, and only the *B. decumbens* cultivar Basilisk was unaffected by high Al^3+^ (Bitencourt *et al.*, 2011). Interestingly, Furlan *et al.* (2018) recently reported a higher Al^3+^ tolerance in *B. brizantha* cv. Xaraes than in *B. decumbens* cv. Basilisk.

The three QTLs we identified were of moderate effect (LOD scores under 6, each accounting 12.8 % to 16.1% of observed phenotypic variance), but were observed for several root traits. DE genes in the QTLs highlighted a link with membrane transport (including metal ions), regulation and signalling (binding to DNA), and energy (carbohydrate metabolism and binding to ATP/ADP/GTP) molecular mechanisms in the three QTLs. The three QTLs were associated with similar molecular functions. Eight genes in the QTL were DE in both species, including two metal-binding transmembrane P450 cytochromes (8440G2 and 598G8), one root-exclusive potassium ion membrane transporter (94G8) induced by potassium starvation (Banuelos *et al.*, 2002), a cysteine XCP1 protease (869G16) that may have a developmental senescence specific cell death function during apoptosis and heavy metal detoxification (De Michele *et al.*, 2009), and a protein (8234G4) not found in other related species.

We are also making publicly available the genome assembly and gene annotation of a diploid accession of *B. ruziziensis*. This diploid accession collected in Burundi is likely the genotype that was mutagenised with colchicine to produce CIAT BRX 44-02, the autotetraploid *B. ruziziensis* progenitor of the interspecific population analysed in this study. Our transcriptomic study and comparative genomics analysis are examples of its utility. While tetraploid *Brachiaria* are the main accessions used for breeding, we opted for a non-polyploid diploid accession because *Brachiaria* grasses are heterozygous outcrossing species.

The assembly was partially scaffolded to the pseudomolecule level using a genetic linkage map of *B. decumbens* CIAT 606 containing approximately 5,000 markers. This genetic map had previously been assembled without a genome (Worthington *et al.*, 2016), but a higher number of markers were incorporated by conducting SNP calling using this new reference. The incorporation of long reads could improve this assembly and allow for polyploid *Brachiaria* genome references in the near future. Because of the limited number of markers, almost all the scaffolds were also placed on the high-quality foxtail millet genome. While we identified three large chromosomal translocations, there was a very high collinearity between the two species as previously observed (Worthington *et al.*, 2016) and as might be predicted from species that we quantified diverged “only” 13-15 MYA.

### Differentially expressed genes associated with internal tolerance mechanisms to Al^3+^ in *Brachiaria*

Aluminium internal tolerance mechanisms involve either modification of the properties of the root cell wall, or the uptake and sequestration of Al^3+^ once it enters the plant (Kochian *et al.*, 2015). Ramos *et al.* (2012) observed that *B. decumbens* accumulated Al^3+^ in the mucilage layer of root apices, which reduced the quantity of Al^3+^ reaching the cell wall and crossing the plasma membrane. Arroyave *et al.* (2013) suggested that the presence of a complex exodermis in *B. decumbens*, absent in *B. ruziziensis*, may contribute to a more efficient exclusion of Al^3+^. On the other hand, a higher concentration of root pectin measured in *B. ruziziensis* during stress could evidence increased apoplastic aluminium binding (Horst *et al.*, 2010).

Changes in the structural properties of cell wall carbohydrates are mediated by expansins, endo-β-1,4-glucanases, xyloglucan transferases and hydrolases (XTH) (e.g. AtXTH31 and AtXTH15) and pectin methylesterases (Kochian *et al.*, 2015; Yang *et al.*, 2011). Contrasting regulation (Up-regulated in one species but down-regulated in the other) in enriched GO terms between *B. decumbens* and *B. ruziziensis*, suggests different regulation of genes associated with xyloglucan transfer (GO:16762), glycosyl hydrolase (MF:16798) and oxidation (GO:52716). This contrasting enrichment was largely due to 12 XTH proteins that were down-regulated during stress in *B. ruziziensis*, but not in *B. decumbens*. One of them (5400G2) was located in the QTL region on LG1 (Chr 8). There were three additional differentially expressed XTH proteins that were up-regulated during stress: 5233G2 (AtXTH23) only in *B. decumbens*, and 6544G2 (AtXTH32) and 11683G4 (AtXTH27) in both *B. decumbens* and *B. ruziziensis*. Related to this, AtSLK2 is involved in cell wall pectin methylesterification in response to Al^3+^ stress (Geng *et al.*, 2017). The *Brachiaria* homologous gene to SLK2 (1984G2) was up-regulated in both *B. ruziziensis* and *B. decumbens*, but only DE under stress in *B. decumbens*.

Once Al^3+^ has entered the root, the uptake and sequestration of Al^3+^ includes molecular binding and eventually compartmentation of the toxin. ALS (Aluminium Sensitive) transporters and NRAM metal ion transporters have been proposed as the key proteins in Al^3+^ localization to the tonoplast and other cell organelles, and away from the sensitive root tips in *A. thaliana* and rice (Huang *et al.*, 2010; Huang *et al.*, 2012; Larsen *et al.*, 2007). Specifically, an NRAM aluminium transporter (NRAT1) localized in the plasma membrane appears to be expressly involved in storing Al^3+^ in root vacuoles in rice and maize. In *Brachiaria*, we identified 21 NRAM proteins, but only three of them were DE: the homologous gene (2499G8) to OsTITANIA (AtOBE3/ATT1), which is the transcription factor that functions as a regulator of NRAM and other metal transporter genes (Tanaka *et al.*, 2018) and two of the three *Brachiaria* proteins showing close homology to NRAT1 (3672G6 and 30107G2). These were significantly up-regulated in both *B. decumbens* and *B. ruziziensis* during stress. Eight genes in the *Brachiaria* genome are homologous to ALS1 and annotated as aluminium-induced ABC transporters, but only one gene (3096G6) was up-regulated during stress in *B. ruziziensis.*

In rice, the complex formed by STAR1 and STAR2/AtALS3 (Sensitive To Aluminium Rhizotoxicity) is involved in aluminium-induced alterations of the cell wall composition related to less aluminium-binding in the apoplast (Huang *et al.*, 2010; Huang *et al.*, 2009). These ABC transporters appear to mediate the efflux of UDP-glucose into the cell wall, which could alter the cell wall composition and lead to a reduction in Al-binding capacity (Kochian *et al.*, 2015). Fifteen proteins were homologous to STAR/ALS in *Brachiaria*, but only two were DE: gene 8448G4 homologous to OsSTAR1, and gene 7361G2 homologous to OsSTAR2. STAR1 and STAR2 were up-regulated in both *B. ruziziensis* and *B. decumbens* during stress.

### Differentially expressed genes associated with external tolerance mechanisms to Al^3+^ in *Brachiaria*

Most aluminium tolerant crops additionally rely on external restriction to prevent the uptake of aluminium and its entry into the root cells through the release of anionic organic acids in the rhizosphere that chelate the Al^3+^ (Kochian *et al.*, 2015; Rao *et al.*, 2016). However, *Brachiaria* appears to not rely on secreted organic acids since there was no difference in secreted organic acids between *Brachiaria* species (Arroyave *et al.*, 2013; Wenzl *et al.*, 2001). The lack of correlation between exudation and resistance has also been observed in rice (Famoso *et al.*, 2010; Ma *et al.*, 2002). Furthermore, tolerant *B. decumbens* accessions secreted 3-30 times less organic acids than sensitive species such as maize and wheat (Arroyave *et al.*, 2018; Wenzl *et al.*, 2001). However, while *B. decumbens* citrate exudation was about 200 times lower than that observed in aluminium-tolerant rice, the same study evidenced high oxalate exudation in *B. decumbens* roots, but only between 24 and 36 hours after exposure to the toxic concentration (Arroyave *et al.*, 2018). It appears that other mechanisms of resistance overshadow the impact of root exudation.

Two families of membrane transporters, aluminium-activated malate transporter (ALMT) and the multidrug and toxic compound extrusion (MATE) family, are responsible for plasma membrane malate and citrate efflux, respectively (Guimaraes *et al.*, 2014; Raman *et al.*, 2005; Rao *et al.*, 2016). Citrate is a much stronger chelating agent for Al^3+^ than malate (Ma, 2000). In *Brachiaria*, we identified three citrate MATE proteins, which were homologous to FRDL (Ferric Reductase Defective Like) proteins and up-regulated with large fold-changes in *B. decumbens* only (61G2), or in both species (259G14 and 675G12). In rice, FRDL4 is responsible for aluminium-induced citrate efflux required for external detoxification (Yokosho *et al.*, 2011).

We identified thirteen aluminium-activated malate transporters (AMLTs) in the genome. However, only two were differentially expressed: 11634G4 was up-regulated 4.74 fold-change during stress in *B. decumbens*, and 462G24 was down-regulated 2.35 fold-change in *B. ruziziensis*. We identified a cluster with three contiguous non-DE AMLTs (5136G2, 5136G4 and 5136G6) in scaffold 5136 (LG3 (Chr. 7): 71.83-72.95 cM) around 10 cM from the QTL. This is consistent with the observation that copy-number variation of ALMT correlated with aluminium resistance in rye and maize (Collins *et al.*, 2008; Maron *et al.*, 2013).

C_2_H_2_-type zinc-finger transcription factors STOP1 (ART1 in rice) and STOP2 regulate aluminium-induced expression of several MATE and ALMT genes in Arabidopsis and rice (Iuchi *et al.*, 2007; Kobayashi *et al.*, 2014; Yamaji *et al.*, 2009). We identified three transcription factors homologous to AtSTOP1, one DE in *B. ruziziensis* (29G2) and another two (3833G12 and 243G34) which were not. We also identified two transcription factors homologous to AtSTOP2, both DE with high fold-change in *B. ruziziensis* and *B. decumbens* (126G26 and 1615G2). All five were homologous to OsART1, the STOP1 homolog in rice, which up-regulated at least 31 genes in an Al-dependent manner, including STAR1, FRDL, NRAMP proteins (Kochian *et al.*, 2015; Yamaji *et al.*, 2009).

The “transmembrane transporters” GO term (MF:22857) were highly overrepresented among up-regulated DE genes during stress in both species (44 genes in *B. decumbens* and 70 genes in *B. ruziziensis*). Fifteen transmembrane transporters were DE in both species, and five were annotated as induced by aluminium: NRAT1 (3672G6), STAR1 (8448G4), ALS3 (7361G2) and two citrate MATE transporter (675G12 and 259G14). All these have been highlighted in our discussion, up-regulated with high fold-changes in both species, and likely indispensable genes for aluminium-tolerance in *Brachiaria*.

In this work we have presented a comprehensive analysis of the molecular mechanism linked to aluminium tolerance in *Brachiaria* species. By assembling and annotating a diploid genotype of *B. ruziziensis* we have developed the capability for genomic-based studies of desirable phenotypic traits. Using this resource, we have identified three QTLs associated to root architecture and vigour during Al^3+^ stress in a hybrid population from a high and low tolerant accession. We have also identified a number of genes and molecular responses that impact on different aspects of signalling, cell-wall composition and active transports as response to aluminium stress. *Brachiaria* tolerance appears to build in the same genes than in rice. However, we found that external mechanisms such sequestration of Al^3+^ common in other grasses might be not that important in *Brachiaria.* Also, contrasting regulation in the same genotype after 8 or 72 hours of Al^3+^ stress of numerous genes involved in RNA translation can explain the different levels of tolerance among different Brachiaria species. The newly annotated draft genome represents an important base upon which study other aspects of *Brachiaria* biology.

## ACKNOWLEDGEMENTS

This work was partially funded by a BBSRC’s Global Challenge Research Fund BB/P028098/1 and a BBSRC’s Newton Fund Postdoctoral Twinning Award BBS/OS/NW/000011.

## DATA AVAILABILITY

Raw reads are deposited in SRA under accession PRJNA437375. The genome assembly is deposit at NCBI with accession GCA_003016355.1; The genome assembly, chromosomal anchoring in AGP format, and gene annotation in GFF3 format can also be downloaded as individual files (http://dx.doi.org/10.5281/zenodo.3703092). Individual scaffolds can be accessed at NCBI’s GenBank accession numbers PVZT01000001 to PVZT01102577.

