## Supplementary Tables and Figures for "A new genome allows the identification of genes associated with natural variation in aluminium tolerance in *Brachiaria* grasses"

- **Supplementary File 1:** Cumulative root length (RL), root biomass (RB), and root tip diameter (RD) during Al^3+^ stress (A) and control (C) conditions, and the ratio (R) between stress and control values, in the interspecific progeny between CIAT 606 and BXR 44-02.
- **Supplementary File 2:** Gene annotation in GFF3 format.
- **Supplementary File 3:** Functional annotation of the genes, including GO terms and homologous proteins in NCBI nr database, Uniprot, *A. thaliana*, rice, *P. halli*, *S. italica* and *S. viridis.*
- **Supplementary File 4:** Assignment of the proteins in the Poaceae family to eggNOG orthologous groups to identify shared clusters of proteins among these species.
- **Supplementary File 5:** Anchoring 21,145 *Brachiaria ruziziensis* scaffolds longer than 10 Kbp or with at least one annotated gene (533.9 Mbp) in *S. italica* nine chromosomes.
- **Supplementary File 6:** Chromosomal position of the 41,974 transcripts in *Brachiaria ruziziensis* based on the synteny with the *S. italica* genome. In BED5 format.
- **Supplementary File 7:** Genetic map with 4,427 markers placed at LOD 10 in 18 linkage groups, including the position of each marker in the genetic map and genome assembly.
- **Supplementary File 8:** Functional annotation of the 84 DE genes within QTLs.
- **Supplementary File 9:** Enrichment analysis of the GO terms (full ontology) over-represented among DE genes in each species with the biological processes (BP) and molecular functions (MF).
- **Supplementary File 10:** Enrichment analysis of the GO SLIM terms (reduced ontology) over-represented among DE genes in each species with the biological processes (BP) and molecular functions (MF).

**Supplementary table S1:** Root length, diameter and biomass in the *B. decumbens* CIAT 606 and *B. ruziziensis* BRX 44-02 (cv. Basilisk) progenitors after growing for 20 days in control and high 200 μM AlCl_3_ concentration hydroponic solutions

|  | **Root Length (mm)** | | | **Root Tip Diameter (mm)** | | | **Root biomass (milligrams)** | | |
| --- | --- | --- | --- | --- | --- | --- | --- | --- | --- |
|  | **Control** | **Stress** | **Ratio** | **Control** | **Stress** | **Ratio** | **Control** | **Stress** | **Ratio** |
| *B. decumbens* CIAT 606 | 428 | 261 | 0.61 | 0.29 | 0.31 | 0.109 | 54 | 34 | 0.63 |
| *B. ruziziensis* BRX 44-02 | 177 | 72 | 0.41 | 0.39 | 0.46 | 0.118 | 30 | 17 | 0.55 |
| Mean population | 474 | 212 | 0.45 | 0.32 | 0.38 | 0.118 | 64 | 39 | 0.61 |

**Supplementary Table S2:** Statistics of the intermediate steps, alternative assemblies, final assembly and pseudo-molecules for the *B. ruziziensis* CIAT 26162 genome.

| Step | Total length (Mbp) | % Ns | Sequences | N50 (Kbp) |
| --- | --- | --- | --- | --- |
| WGS Platanus | 712.4 | 17.45 | 196,321 | 17.4 |
| ABySS+SOAP2 (Discarded) | 815.4 | 12.74 | 268,486 | 5.5 |
| Pacbio Gapfilling | 796.8 | 11.39 | 191,540 | 23.3 |
| Deposit WGS (GCA_003016355) | 732.5 | 10.59 | 102,579 | 27.8 |
| Unanchored reference (Sequences over 10Kb) | 533.9 | 11.7 | 23,076 | 44.6 |
| Anchored in 9 chrs | 525.1 | 12.18 | 9 | 55.88*Mbp |

**Supplementary Table S3: Classification of the repeat content in the Brachiaria genome.**

| **Category** | **Superfamily** | **Coverage (bps)** | **Fraction genome*** |
| --- | --- | --- | --- |
| Class 1 Transposable elements (TEs) | Gypsy | 156,824,480 | 23.9 |
|  | Copia | 62,486,851 | 9.5 |
|  | Pao | 55,071 | 0.0 |
|  | Other LTRs | 972,934 | 0.1 |
|  | SINEs | 2,939,316 | 0.4 |
|  | LINEs | 11,929,645 | 1.8 |
|  |  | (235,208,297) | (35.8) |
| Class 2 (DNA) Transposable elements (TEs) | hAT | 631,797 | 0.1 |
|  | hAT_Ac | 2,534,322 | 0.4 |
|  | hAT_Tag1 | 1,082,474 | 0.2 |
|  | hAT_Tip100 | 306,963 | 0.0 |
|  | Harbinger/PIF | 9,477,600 | 1.4 |
|  | MULE | 7,895,375 | 1.2 |
|  | Stowaway | 4,436,872 | 0.7 |
|  | CMC_EnSpm | 27,724,058 | 4.2 |
|  | Helitron | 1,339,398 | 0.2 |
|  |  | (55,428,859) | (8.4) |
| Non TEs | Unclassified TE | 40,000,354 | 6.1 |
|  | Simple Repeats | 482,035 | 0.1 |
|  | Satellites | 2,999,591 | 0.5 |
|  |  | (43,481,980) | (6.6) |
| Unclassified TE | Other | 335,155 | 0.1 |
| **TOTAL** |  | **334,454,291** | **51.0** |

*656Mbp after excluding ambiguous nucleotides (Ns)

**Supplementary Table S4:** Alignment of the transcripts and proteins from five sequenced species in the Panicoideae subfamily [foxtail millet (*Setaria italica*), green foxtail (*Setaria viridis* (L.) Beauv.), *Panicum halli* Vasey, switchgrass (*Panicum virgatum* L.), and maize (*Zea mays* L.)], in the *Brachiaria ruziziensis* genome with a minimum identify of 70 %. Transcripts (longest one per gene) were aligned with GMAP and proteins were aligned with Exonerate. Sequences were obtained from Phytozome v.12 or Ensembl (v.284) in the case of maize.

|  | **Species** | **Total** | **PID>70%** | | **PID>70% & PCOV>50%** | |
| --- | --- | --- | --- | --- | --- | --- |
| **Transcripts** | ***S. italica*** | 43,001 | 37,449 | 87.1 | 29,534 | 68.7 |
|  | ***S. viridis*** | 47,205 | 36,372 | 77.1 | 23,110 | 49 |
|  | ***P. halli*** | 49,852 | 40,818 | 81.9 | 31,599 | 63.4 |
|  | ***P. virgatum*** | 91,838 |  |  |  |  |
|  | ***Z. mays*** | 88,760 | 58,312 | 65.7 | 36,642 | 41.3 |
| **Proteins** | ***S. italica*** | 43,001 | 34,749 | 80.8 | 29,975 | 69.7 |
|  | ***S. viridis*** | 47,205 | 33,157 | 70.2 | 27,953 | 59.2 |
|  | ***P. halli*** | 49,852 | 37,516 | 75.3 | 32,753 | 65.7 |
|  | ***P. virgatum*** | 91,838 |  |  |  |  |
|  | ***Z. mays*** | 88,760 | 54,091 | 60.9 | 45,951 | 51.8 |

**Supplementary Table S5:** EggNOG clusters in six sequenced species in the Panicoideae subfamily, *B. ruziziensis*, foxtail millet (*Setaria italica*), green foxtail (*Setaria viridis* (L.) Beauv.), *Panicum halli* Vasey, switchgrass (*Panicum virgatum* L.), and maize (*Zea mays* L.), classified by number of proteins per cluster.

|  | ***P. virgatum*** | | ***S. italica*** | | ***S.viridis*** | | ***P. halli*** | | ***B. ruziziensis*** | | ***Z. mays*** | |
| --- | --- | --- | --- | --- | --- | --- | --- | --- | --- | --- | --- | --- |
| **GENES** | **Num** | **%** | **Num** | **%** | **Num** | **%** | **Num** | **%** | **Num** | **%** | **Num** | **%** |
| **1** | 1897 | 8.4 | 18629 | 84.4 | 18508 | 83.4 | 18413 | 86.2 | 12572 | 66.7 | 14369 | 70.3 |
| **2** | 11955 | 53.1 | 2432 | 11.0 | 2534 | 11.4 | 2176 | 10.2 | 3908 | 20.7 | 4250 | 20.8 |
| **3** | 4602 | 20.5 | 559 | 2.5 | 630 | 2.8 | 464 | 2.2 | 1164 | 6.2 | 1040 | 5.1 |
| **4** | 1808 | 8.0 | 193 | 0.9 | 235 | 1.1 | 157 | 0.7 | 472 | 2.5 | 381 | 1.9 |
| **>4** | 2237 | 9.9 | 263 | 1.2 | 285 | 1.3 | 163 | 0.8 | 733 | 3.9 | 411 | 2.0 |
| **total** | 22499 |  | 22076 |  | 22192 |  | 21373 |  | 18849 |  | 20451 |  |

**Supplementary Table S6:** Peak and interval positions for the identified QTLs, as well as corresponding *S. italica* chromosome.

| **Trait*** | **LG** | **Peak marker** | **Peak Position (cM)** | **Position interval (cM)** | **Marker interval** | **LOD** | **R2** | **additive effect** | **Si*** |
| --- | --- | --- | --- | --- | --- | --- | --- | --- | --- |
| RLA | 1 | scaf_70183_123 | 12.65 | 5.22 - 31.250 | scaf_245_124360 - scaf_1729_44015 | 4.81 | 13.6 | -22.31 | 8 |
| RLC | 1 | scaf_2065_28271 | 26.027 | 5.22 - 28.893 | scaf_1787_41196 - scaf_3809_36896 | 5.78 | 16.1 | -45.05 | 8 |
| RRL | 3 | scaf_1152_23964 | 96.802 | 88.62-98.851 | scaf_2718_14282 - scaf_5425_26287 | 4.75 | 13.4 | 3.12 | 7 |
| RBA | 1 | scaf_1948_47501 | 5.22 | 5.22 - 32.481 | scaf_245_124360 - scaf_1218_39495 | 5.14 | 14.4 | -0.003 | 8 |
| RBC | 1 | scaf_18010_6509 | 25.797 | 17.162 - 28.893 | scaf_7830_644 - scaf_3809_36896 | 5.25 | 14.7 | -0.004 | 8 |
| RRD | 3 | scaf_14238_7306 | 79.738 | 79.738 - 83.853 | scaf_14238_7306 - scaf_298_25199 | 4.02 | 11.5 | -2.36 | 7 |
| RRD | 4 | scaf_11042_5202 | 50.423 | 49.12 - 62.127 | scaf_1413_25183 - scaf_1181_29646 | 4.54 | 12.8 | 2.48 | 3 |

*Si: *Setaria italica* chromosome.

RLA: Root length in Al^3+^ stress; RLC: Root length in control; RRD: Relative root length ratio (stress/control); RB: Root biomass; RD: Root tip diameter.

**Supplementary Table S7:** Enrichment analysis of the GO SLIM terms over-represented among DE genes in B. ruziziensis BRX 44-02 (Bruz), B. decumbens CIAT 606 (cv. Basilisk) (Bdec), or PRJNA314352 from Salgado *et al*. (2017).

| **GO term** | **MOLEC. FUNC.** | **Bdec CIAT 606** | | | **Bruz BRX 44-02** | | | **Salgado *et al,* 2017** | | |
| --- | --- | --- | --- | --- | --- | --- | --- | --- | --- | --- |
|  |  | **Pval** | **REG** | **GENES** | **Pval** | **REG** | **GENES** | **Pval** | **REG** | **GENES** |
| GO:0003723 | RNA binding (3723) | 0.05709 | down | 14 | 0.40064 | down | 40 | 0.988 | down | 3 |
| GO:0003729 | mRNA binding (3729) | 0.13999 | down | 2 | 0.00358 | down | 8 | 0.419 | down | 1 |
| GO:0003735 | structural constituent of ribosome (3735) | 5.5E-12 | down | 24 | 1E-30 | down | 112 | 0.93632 | up | 5 |
| GO:0005198 | structural molecule activity (5198) | 0.19102 | up | 4 | 0.02756 | down | 120 | 0.583 | down | 2 |
| GO:0008092 | cytoskeletal protein binding (8092) | 0.58493 | down | 1 | 0.25288 | down | 5 | 0.035 | down | 3 |
| GO:0008134 | transcription factor binding (8134) | 0.0312 | up | 3 | 0.85829 | down | 1 | 1 | 0 | 0 |
| GO:0008289 | lipid binding (8289) | 0.09253 | up | 5 | 0.21271 | down | 9 | 0.26684 | up | 5 |
| GO:0008565 | protein transporter activity (8565) | 0.08093 | up | 3 | 0.14489 | down | 5 | 0.446 | down | 1 |
| GO:0016491 | oxidoreductase activity (16491) | 0.00034 | down | 40 | 6.8E-09 | down | 143 | 0.00081 | up | 69 |
| GO:0016757 | glycosil transferase (16757) | 0.63379 | up | 7 | 0.0438 | up | 22 | 0.20773 | up | 14 |
| GO:0016765 | alkyl transferase (16765) | 0.46513 | down | 3 | 0.03894 | down | 16 | 0.000019 | up | 17 |
| GO:0016798 | glycosyl hydrolase (16798) | 0.00084 | up | 18 | 0.0002 | down | 39 | 0.00041 | up | 24 |
| GO:0016829 | lyase activity (16829) | 0.01422 | up | 10 | 0.16411 | down | 16 | 2E-12 | up | 30 |
| GO:0016853 | isomerase activity (16853) | 0.43965 | down | 4 | 0.00077 | down | 26 | 0.302 | down | 4 |
| GO:0016874 | ligase activity (16874) | 0.11268 | up | 8 | 0.1587 | up | 13 | 0.0492 | up | 12 |
| GO:0019843 | rRNA binding (19843) | 0.04791 | down | 3 | 0.0000021 | down | 14 | 0.81806 | up | 1 |
| GO:0019899 | enzyme binding (19899) | 0.00567 | down | 8 | 0.90566 | down | 7 | 0.65 | down | 2 |
| GO:0022857 | transmembrane transporter activity (22857) | 0.00000018 | up | 44 | 0.00000031 | up | 70 | 0.00000015 | up | 57 |
| GO:0030234 | enzyme regulator activity (30234) | 0.58307 | down | 3 | 0.00354 | down | 22 | 0.09403 | up | 10 |
| GO:0030674 | protein binding bridging (30674) | 1 | 0 | 0 | 0.06958 | down | 2 | 1 | 0 | 0 |
| GO:0043167 | ion binding (43167) | 0.23666 | up | 44 | 0.0042 | up | 100 | 0.261 | down | 26 |
| GO:0051082 | unfolded protein binding (51082) | 0.09328 | down | 2 | 0.32559 | down | 3 | 1 | 0 | 0 |

REG: Either up-regulated (up) or down-regulated (down)

*Supplementary Table S7 -Cont.-*

| **GO term** | **BIOLOG. PROCESS.** | **Bdec CIAT 606** | | | **Bruz BRX 44-02** | | | **Salgado *et al,* 2017** | | |
| --- | --- | --- | --- | --- | --- | --- | --- | --- | --- | --- |
|  |  | **Pval** | **REG** | **GENES** | **Pval** | **REG** | **GENES** | **Pval** | **REG** | **GENES** |
| **GO:0005975** | carbohydrate metabolic process (5975) | 0.000087 | 1 | 21 | 0.00518 | -1 | 42 | 0.00193 | 1 | 24 |
| **GO:0006091** | generation of precursor metabolites (6091) | 0.44019 | -1 | 4 | 0.0175 | 1 | 15 | 0.0000086 | -1 | 13 |
| **GO:0006397** | mRNA processing (6397) | 0.06748 | -1 | 5 | 0.6105 | 1 | 5 | 0.91865 | 1 | 2 |
| **GO:0006412** | translation (6412) | 0.0002 | -1 | 18 | 1E-30 | -1 | 103 | 0.90928 | 1 | 9 |
| **GO:0006457** | protein folding (6457) | 0.17529 | -1 | 3 | 0.01588 | -1 | 12 | 0.7229 | -1 | 1 |
| **GO:0006464** | cellular protein modification process (6464) | 0.03977 | -1 | 23 | 0.9702 | 1 | 28 | 0.3059 | -1 | 16 |
| **GO:0006520** | cellular amino acid metabolic process (6520) | 0.4388 | 1 | 6 | 0.4677 | 1 | 11 | 0.03173 | 1 | 14 |
| **GO:0006629** | lipid metabolic process (6629) | 0.00645 | -1 | 14 | 0.00613 | -1 | 41 | 0.02375 | 1 | 20 |
| **GO:0006810** | transport (6810) | 0.0114 | 1 | 35 | 0.03057 | -1 | 83 | 0.6354 | -1 | 12 |
| **GO:0006913** | nucleocytoplasmic transport (6913) | 0.0052 | 1 | 7 | 0.232 | 1 | 6 | 0.7814 | -1 | 1 |
| **GO:0006914** | autophagy (6914) | 1 | -1 | 0 | 0.1436 | 1 | 3 | 0.0812 | 1 | 3 |
| **GO:0006950** | response to stress (6950) | 0.591 | 1 | 14 | 0.4765 | 1 | 29 | 0.0521 | -1 | 16 |
| **GO:0007005** | mitochondrion organization (7005) | 0.08128 | -1 | 3 | 0.02374 | -1 | 9 | 0.57158 | 1 | 2 |
| **GO:0007010** | cytoskeleton organization (7010) | 0.76433 | -1 | 1 | 0.13705 | -1 | 9 | 0.0371 | -1 | 4 |
| **GO:0007155** | cell adhesion (7155) | 1 | -1 | 0 | 0.0699 | 1 | 1 | 1 | -1 | 0 |
| **GO:0007165** | signal transduction (7165) | 0.1634 | 1 | 15 | 0.0845 | 1 | 29 | 0.09511 | 1 | 23 |
| **GO:0009056** | catabolic process (9056) | 0.13519 | -1 | 17 | 0.20889 | -1 | 58 | 1.2E-09 | 1 | 58 |
| **GO:0009058** | biosynthetic process (9058) | 0.09869 | -1 | 52 | 0.0267 | 1 | 95 | 0.1459 | -1 | 35 |
| **GO:0019748** | secondary metabolic process (19748) | 0.011 | 1 | 13 | 0.0066 | 1 | 22 | 0.00064 | 1 | 21 |
| **GO:0022618** | ribonucleoprotein complex assembly (22618) | 0.00041 | -1 | 8 | 0.000012 | -1 | 21 | 0.85132 | 1 | 2 |
| **GO:0030154** | cell differentiation (30154) | 0.5706 | 1 | 1 | 0.05932 | -1 | 6 | 0.4517 | -1 | 1 |
| **GO:0030198** | extracellular matrix organization (30198) | 1 | -1 | 0 | 0.0699 | 1 | 1 | 0.05441 | 1 | 1 |
| **GO:0042592** | homeostatic process (42592) | 0.3365 | 1 | 7 | 0.0526 | 1 | 17 | 0.00568 | 1 | 17 |
| **GO:0044281** | small molecule metabolic process (44281) | 0.04068 | -1 | 19 | 0.00017 | -1 | 75 | 3.6E-09 | 1 | 60 |
| **GO:0048856** | anatomical structure development (48856) | 0.4292 | 1 | 13 | 0.0384 | 1 | 29 | 0.8382 | -1 | 7 |
| **GO:0051186** | cofactor metabolic process (51186) | 0.1532 | -1 | 6 | 0.01469 | -1 | 24 | 0.000004 | 1 | 21 |
| **GO:0051301** | cell division (51301) | 1 | -1 | 0 | 0.00458 | -1 | 7 | 1 | 0 | 0 |
| **GO:0055085** | transmembrane transport (55085) | 0.14685 | -1 | 3 | 0.30994 | -1 | 7 | 0.01203 | 1 | 7 |
| **GO:0071554** | cell wall organization or biogenesis (71554) | 0.0000041 | 1 | 17 | 2.8E-14 | -1 | 51 | 0.01151 | 1 | 14 |

**Supplementary Figure S1:** 31mer frequency analysis comparing the short-reads assemblies produced with *Platanus assembler* or the alternative approach using the combination of ABySS for isotigs assembly and SOAP2 for scaffolding. The area under the curve of the Kmer spectra has been coloured according to the number of times that such K-mers appear in the assembly: none in back, once in red, twice in orange, etc.

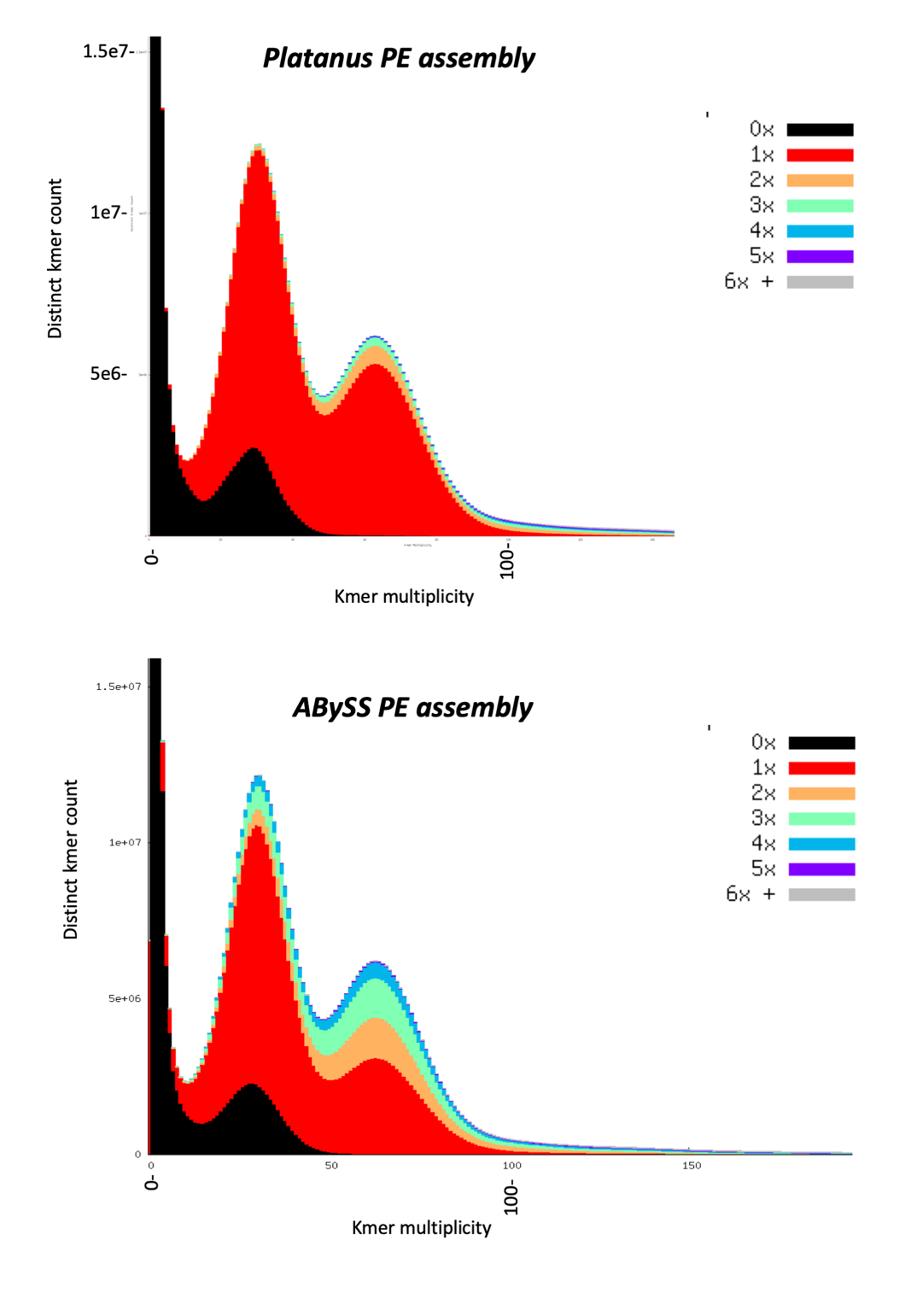

**Supplementary Figure S2:** Divergence (Kimura) rates between the flanking tails in each *Gypsy* and *Copia* LTR duplication events in the *Brachiaria* genome.

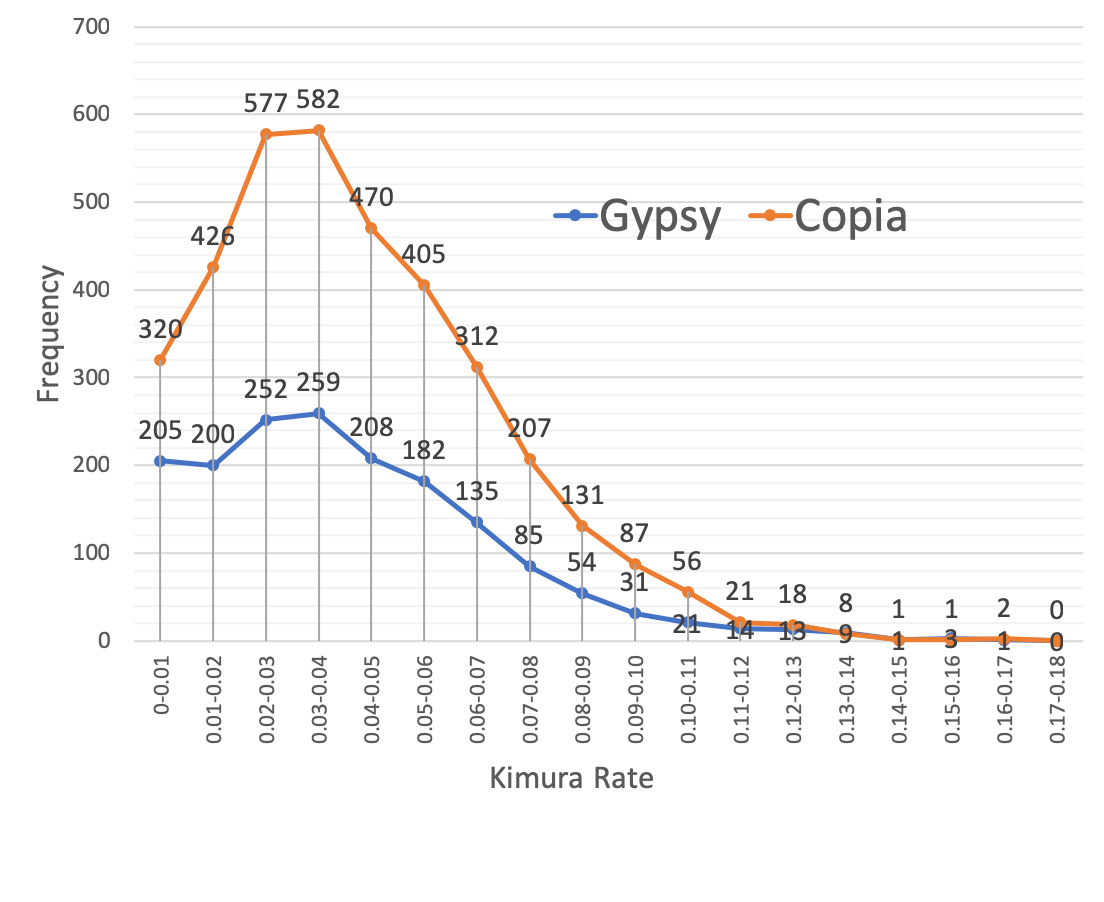

**Supplementary Figure S3:** Species of the top Blastp hit for each 35,982 of the coding transcripts which had a homologous protein in the NCBI non-redundant (nr) database.

Number of best BLAST aligments

|  | Number of best alignments |
| --- | --- |
| Setaria italica | 21109 |
| Dichanthelium oligosanthes | 4610 |
| Sorghum bicolor | 3900 |
| Zea mays | 2229 |
| Oryza sativa Japonica Group | 1684 |
| Oryza sativa Indica Group | 484 |
| Brachypodium distachyon | 399 |
| Aegilops tauschii | 291 |
| Oryza brachyantha | 266 |
| Triticum urartu | 211 |
| Hordeum vulgare subsp. vulgare | 101 |
| Saccharum hybrid cultivar R570 | 98 |
| Triticum aestivum | 47 |
| Phyllostachys edulis | 43 |
| Daucus carota subsp. sativus | 23 |
| Vitis vinifera | 23 |
| Panicum virgatum | 22 |
| Ananas comosus | 21 |
| Oryza sativa | 21 |
| Cephalotus follicularis | 15 |
| Phoenix dactylifera | 15 |
| Elaeis guineensis | 15 |
| Cajanus cajan | 14 |
| Tinamus guttatus | 14 |
| Oryza australiensis | 13 |
| Populus trichocarpa | 13 |
| Saccharum officinarum | 10 |
| Hordeum vulgare | 10 |
| Zea mays subsp. mays | 10 |
| Musa acuminata subsp. malaccensis | 10 |

**Supplementary Figure S4:** Shared eggnog clusters of proteins among *Brachiaria ruziziensis* (Bruz)*,* foxtail millet [*S. italica* (Sita)], *S. viridis* (Svir), maize [*Z. mays* (Zmays)], *Panicum halli* (Phal) and switchgrass [*P. virgatum* (Pvir)]. The “UpSet” plot format provides an efficient way to visualize the intersections (columns) of six species (Rows).

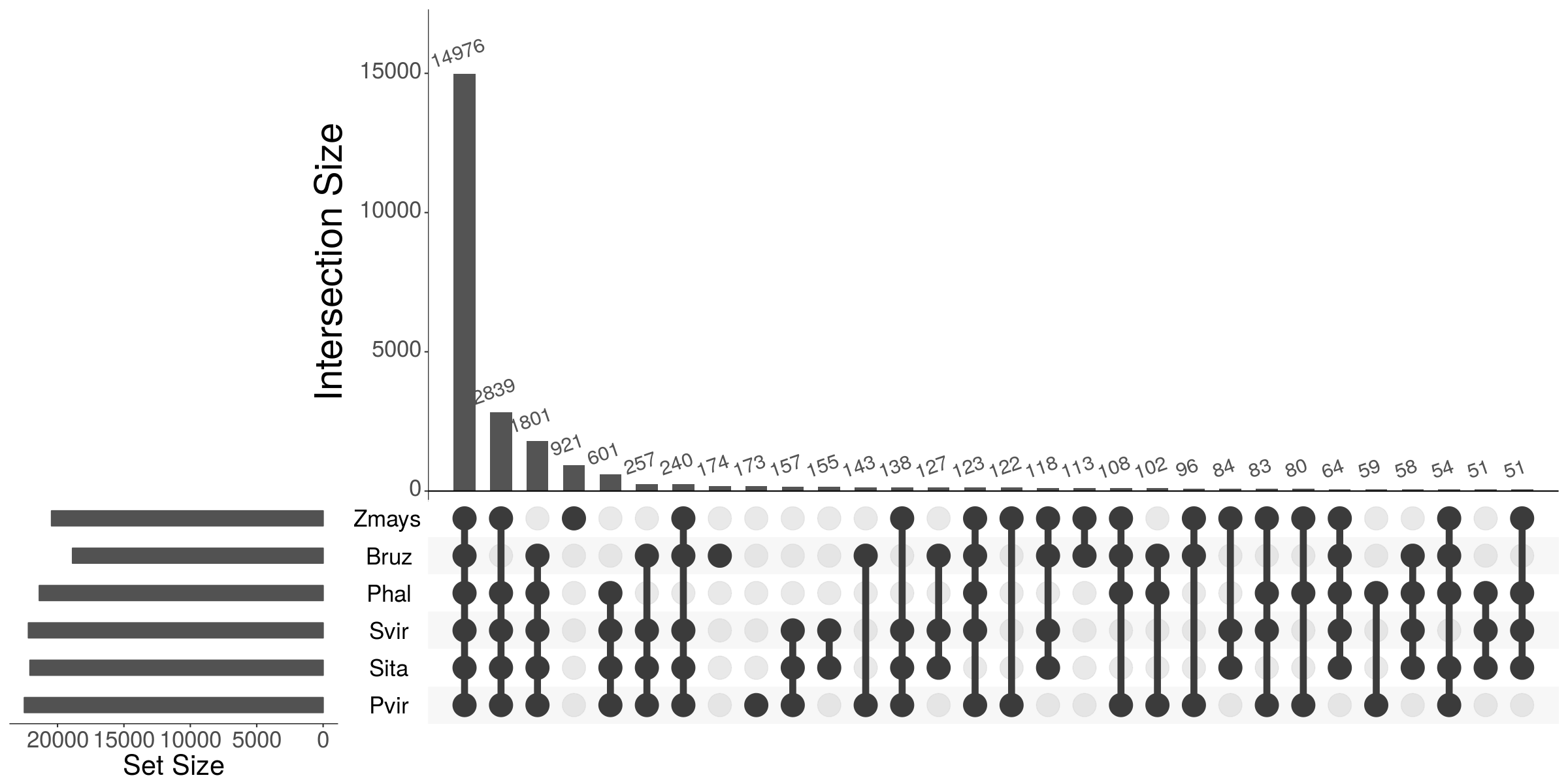

**Supplementary Figure S5:** Kimura rates between homologous gene pairs between *B. ruziziensis* and sequenced relatives including foxtail millet [*S. italica* (Sita)], *S. viridis* (Svir), maize [*Z. mays* (Zmays)], and *P. halli* (Phal)]. Gene pairs were build based on eggNOG clusters.

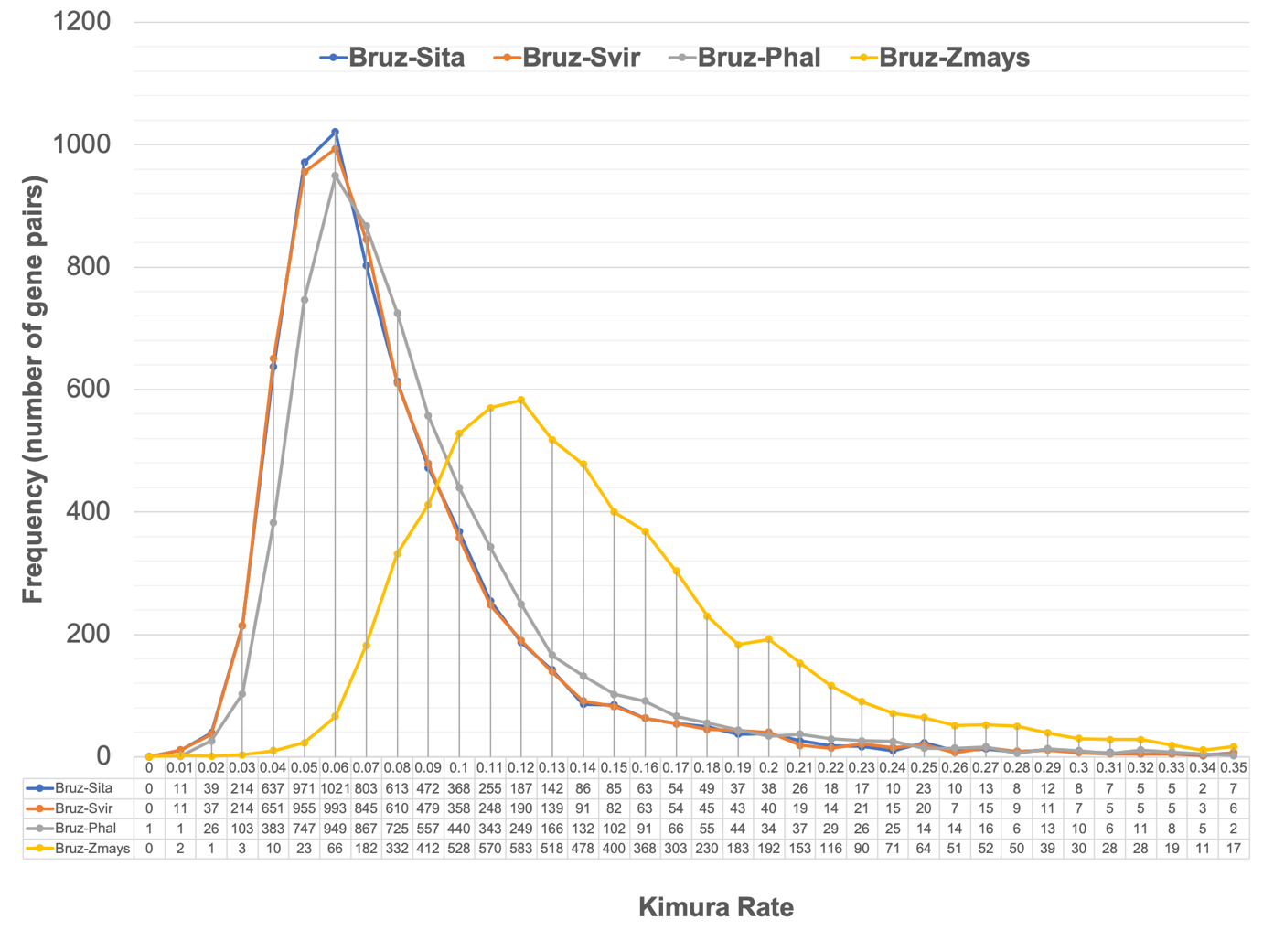

**Supplementary Figure S6:** Phylogenetic tree based on nucleotide divergence rate between sequences in the same eggnog cluster from *B. ruziziensis* and sequenced relatives including foxtail millet [*S. italica* (Sita)], *S. viridis* (Svir), maize [*Z. mays* (Zmays)], and *P. halli* (Phal)].

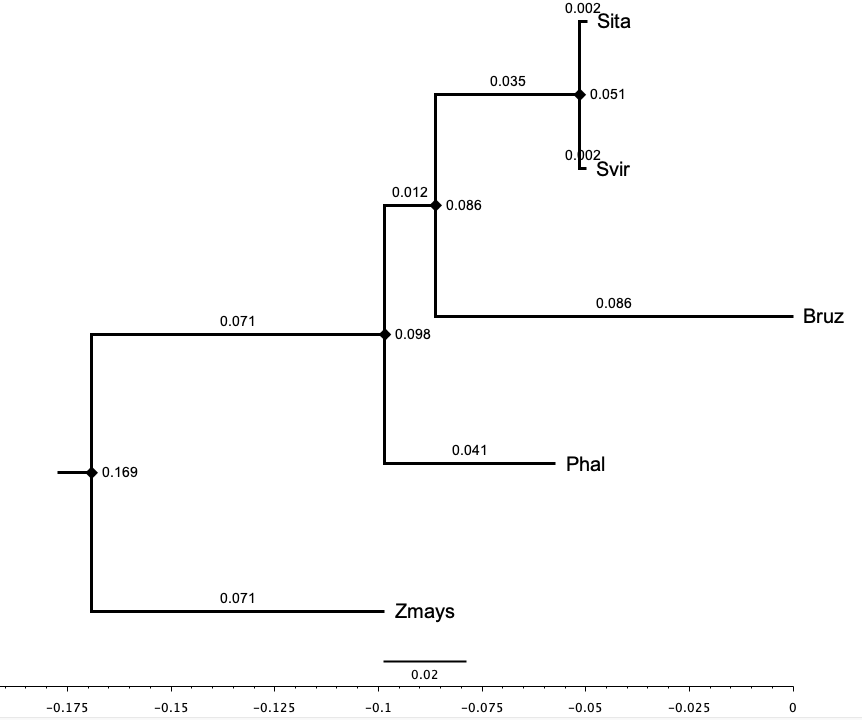

**Supplementary Figure S7:** The final genetic map for the B. decumbens CIAT 606 (cv. Basilisk) progenitor of the interspecific population included 4,427 markers placed at LOD 10 in 18 linkage groups

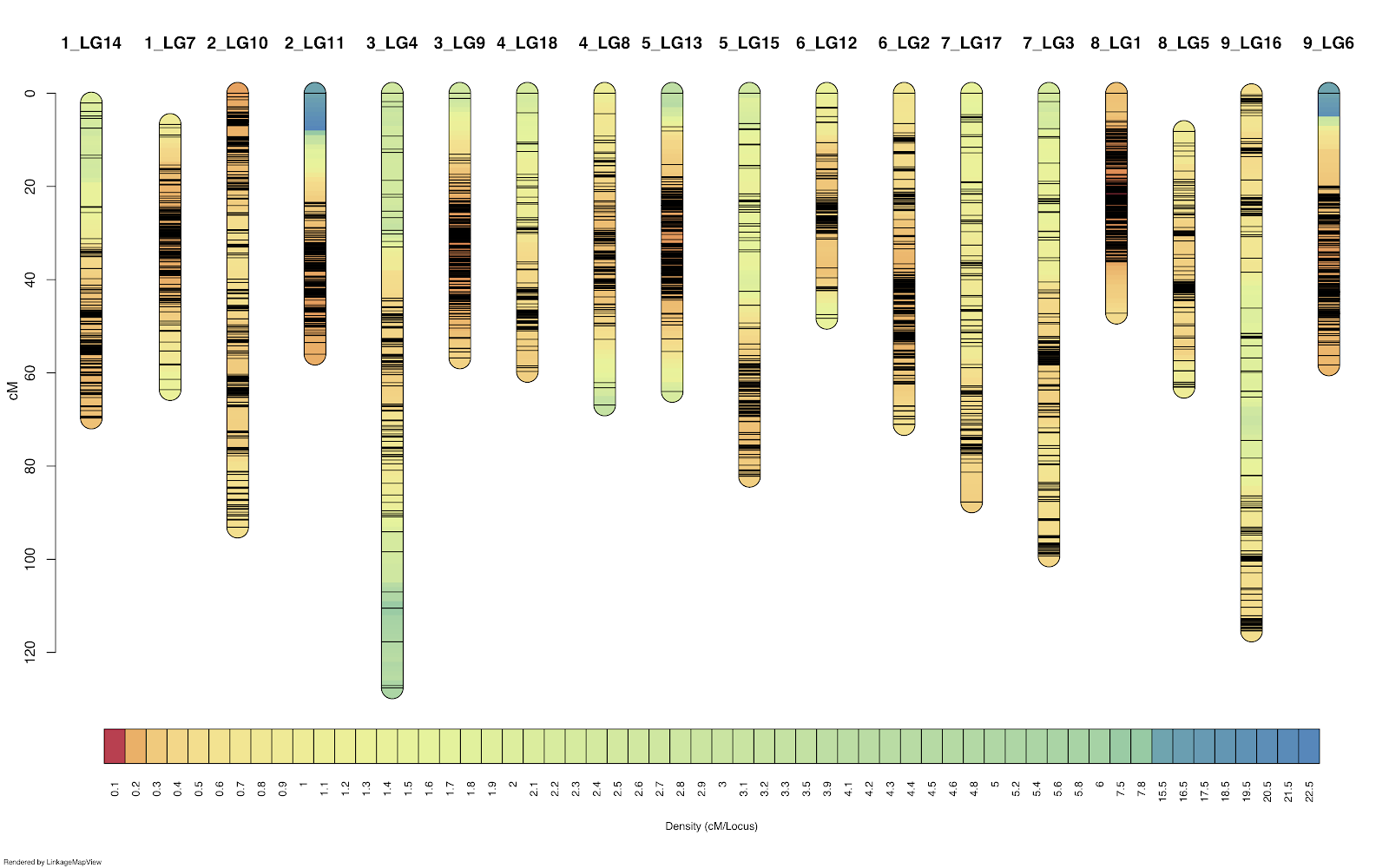

**Supplementary Figure S8:** RNA-seq from stem and root tissue samples extracted from the *B. decumbens* and *B. ruziziensis* progenitors. We also incorporated a reference-based reanalysis of public RNA-seq data (PRJNA314352) from *B. decumbens* var. Basilisks roots (Salgado et al., 2017, Plant Growth Regulation, 83,1:157-170). When the normalised counts for all the genes were used to cluster the samples, these clusters firstly grouped by tissue, secondly by genotype, and thirdly by treatment.

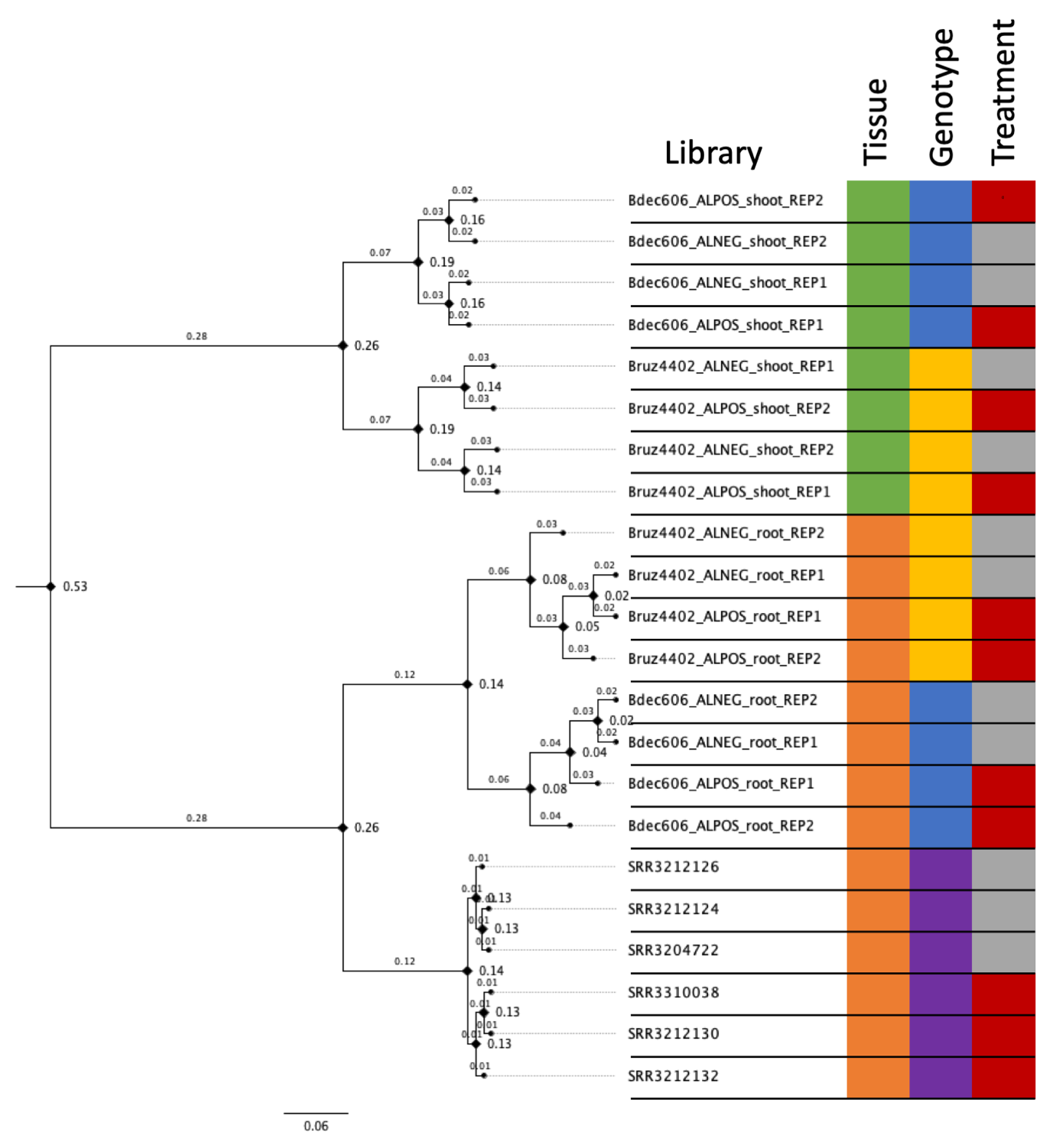

**Supplementary Figure S9:** Enrichment analysis of the “Molecular function” GO terms overrepresented among differentially expressed upregulated (red) or downregulated (blue) genes in roots in *B. decumbens* CIAT 606 and *B. ruziziensis* BRX 44-02.

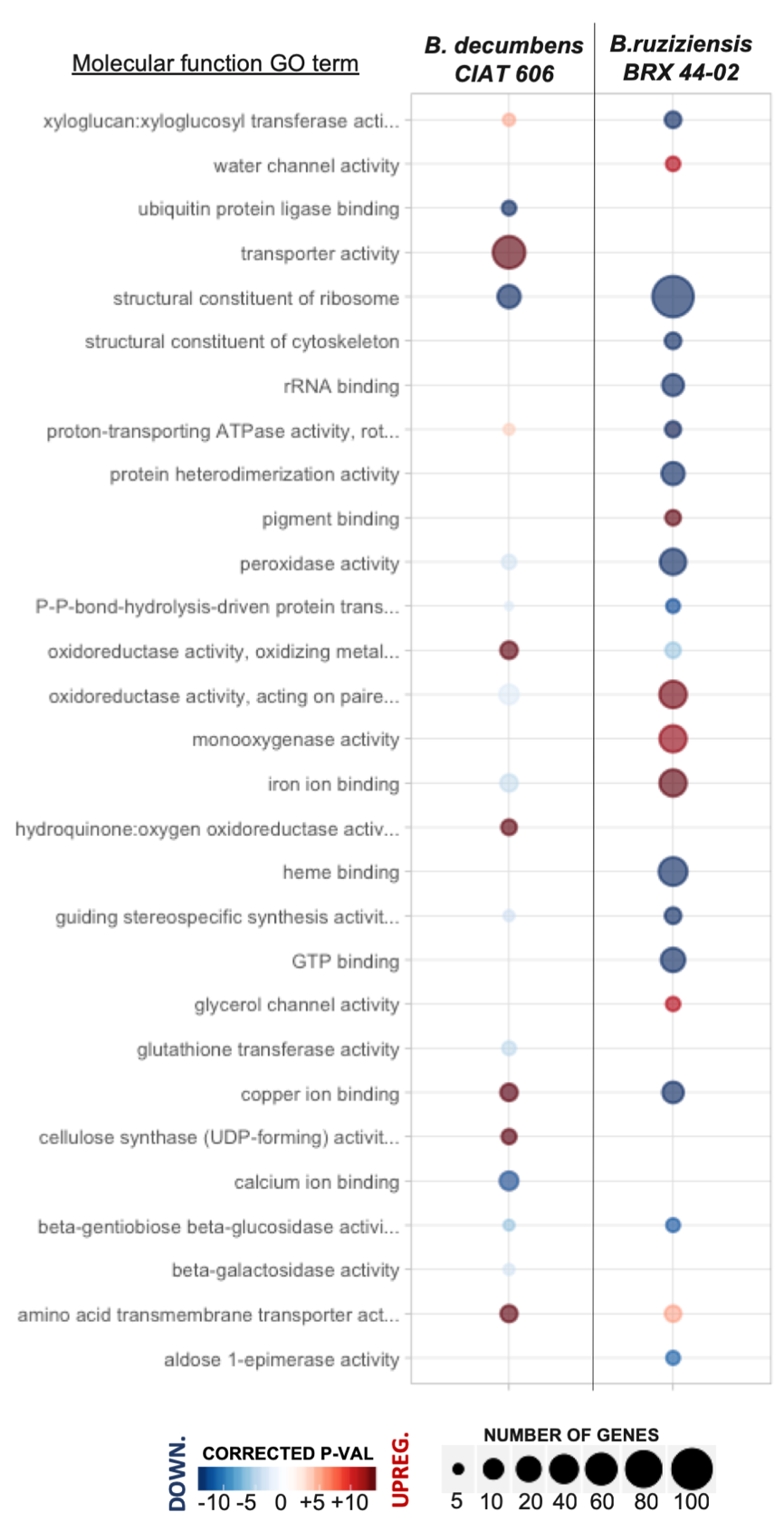

**Supplementary Figure S10:** Enrichment analysis of the “Biological Process” GO terms overrepresented among differentially expressed upregulated (red) or downregulated (blue) genes in roots in *B. decumbens* CIAT 606 and *B. ruziziensis* BRX 44-02.

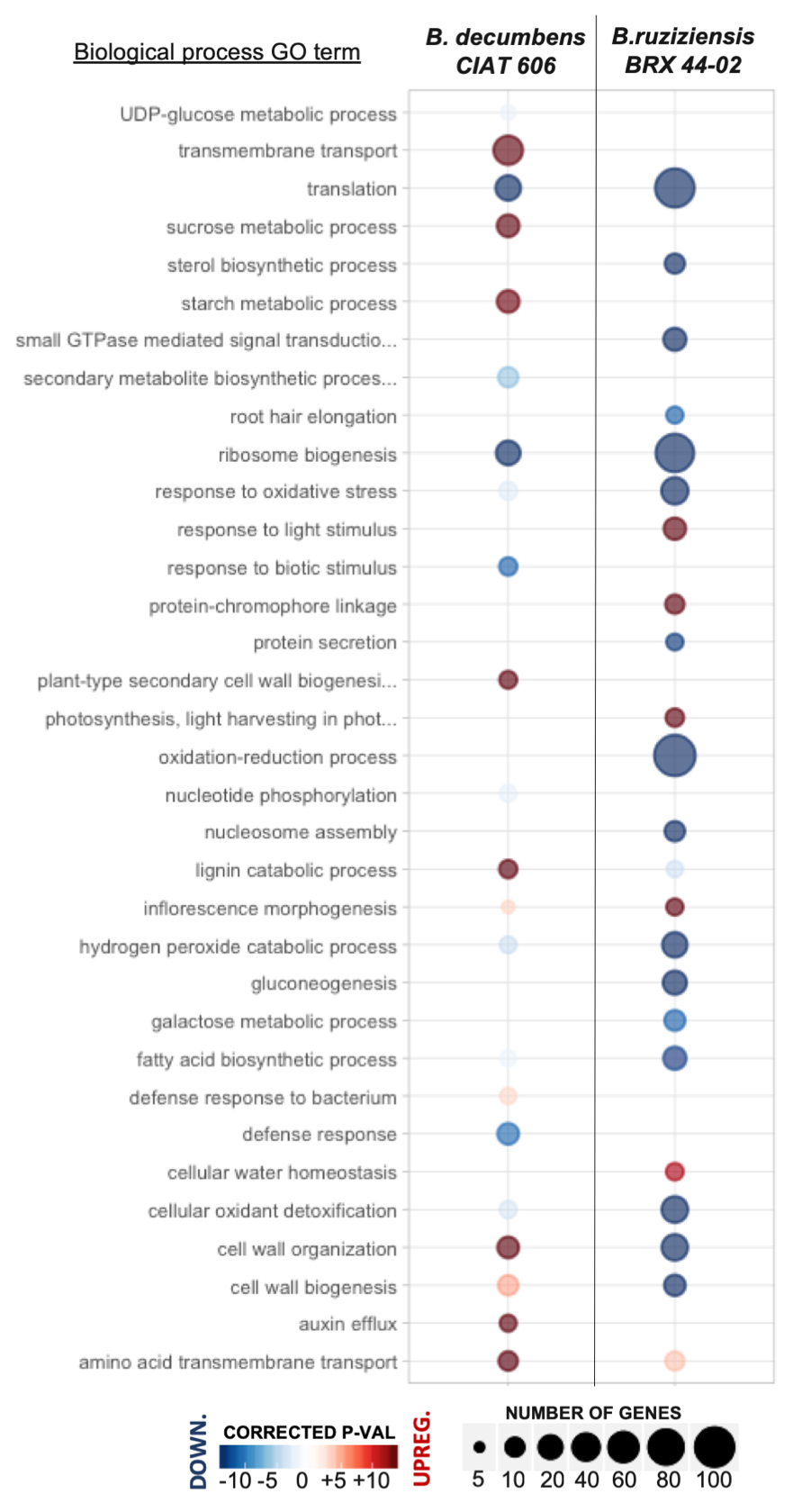

**Supplementary Figure S11:** Correlation matrix plot among GO terms based on the DE genes included in each annotation.

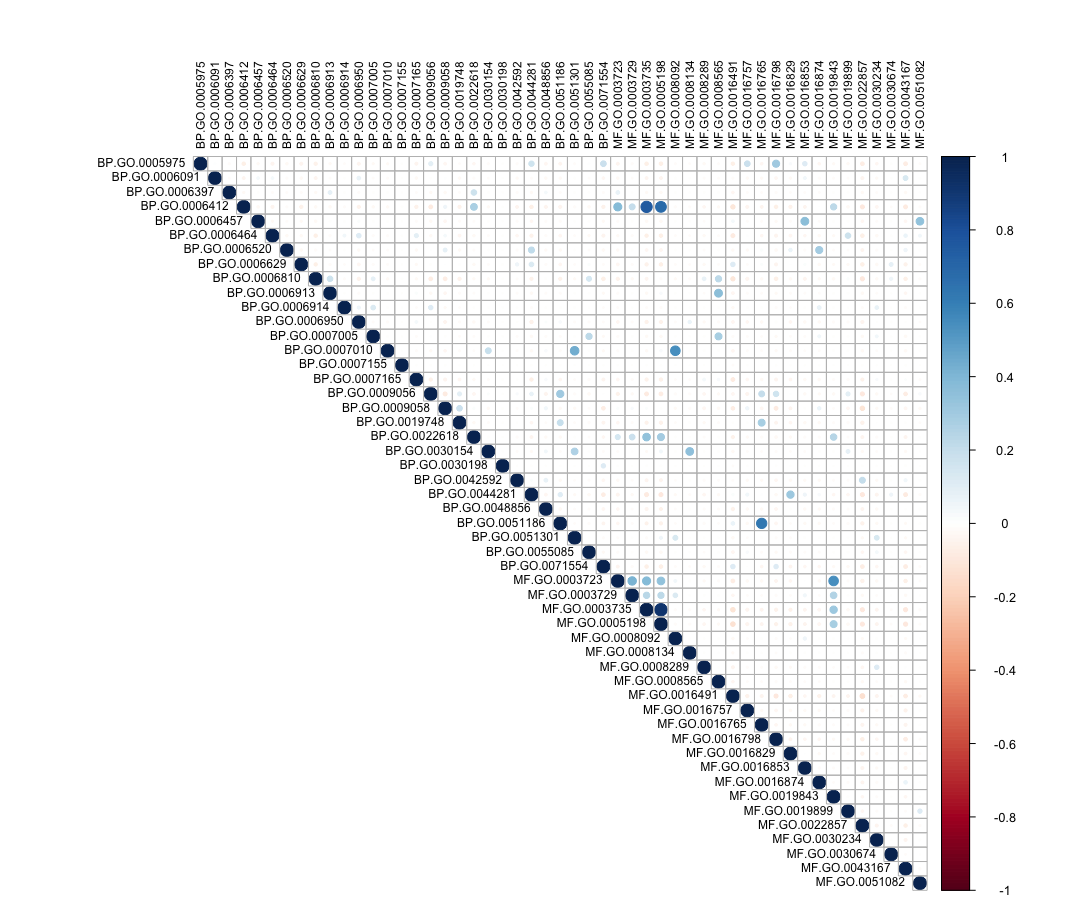

**Supplementary Figure S12:** Comparison the enriched GO Slim terms between *B. decumbens* cv. Basilisk exposed to 200 μM AlCl_3_ for 72 hours and 8 hours, the latter from the reanalysis of public raw data from Salgado *et al*. 2017.

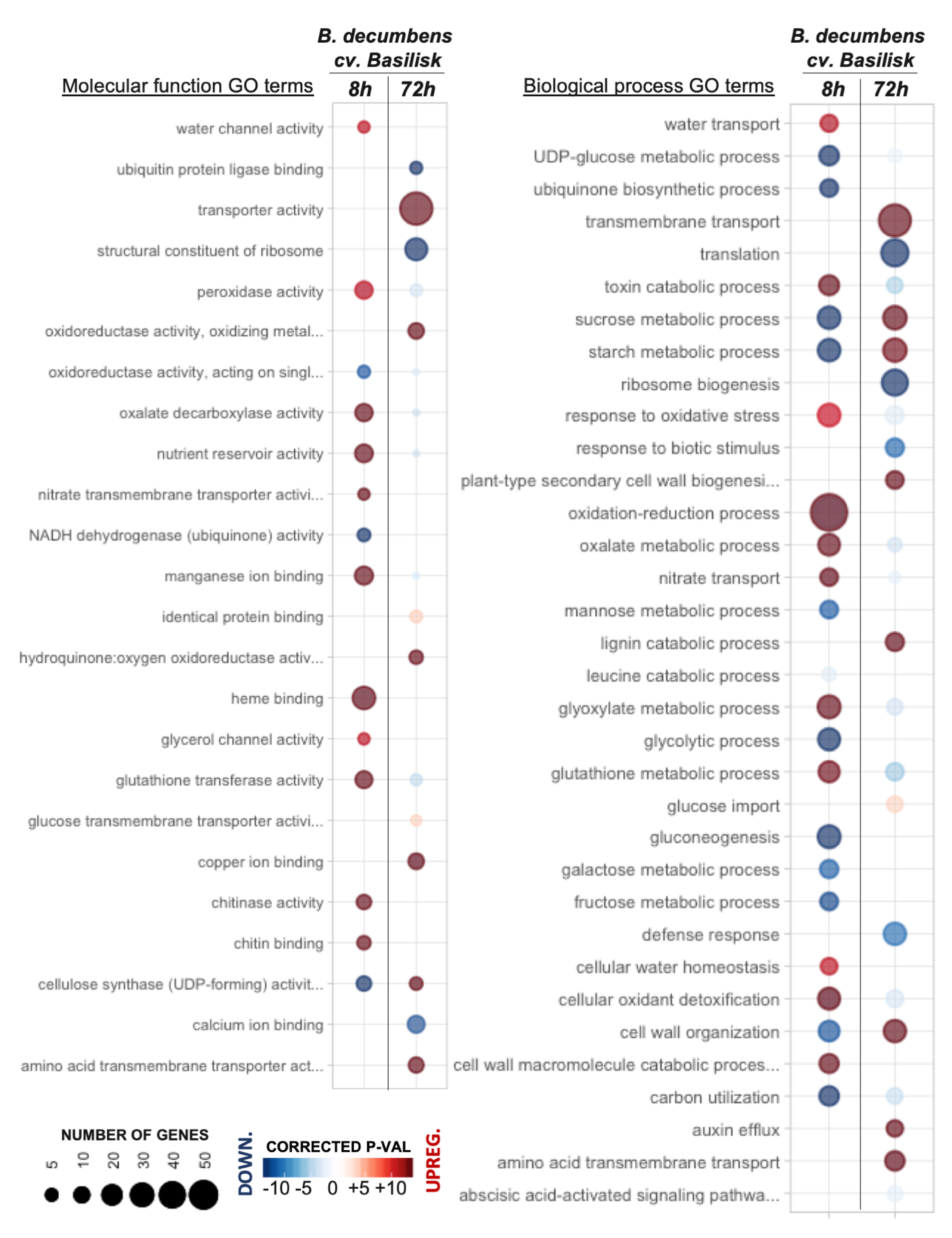
